# Prenatal VEGF Nano-Delivery Reverses Congenital Diaphragmatic Hernia-Associated Pulmonary Abnormalities

**DOI:** 10.1101/2024.02.20.581170

**Authors:** Stavros P. Loukogeorgakis, Federica Michielin, Noura Al-Juffali, Julio Jimenez, Soichi Shibuya, Jessica Allen-Hyttinen, Patrice Eastwood, Ahmed S.N. Alhendi, Joseph Davidson, Eleonora Naldi, Panagiotis Maghsoudlou, Alfonso Tedeschi, Sahira Khalaf, Aziza Khabbush, Manuela Plate, Camila Fachin, Andre Dos Santos Dias, Nikhil Sindhwani, Dominic Scaglioni, Theodoros Xenakis, Neil Sebire, Monica Giomo, Simon Eaton, Jaan Toelen, Camilla Luni, Piero Pavan, Peter Carmeliet, Francesca Russo, Samuel Janes, Marko Z. Nikolic, Nicola Elvassore, Jan Deprest, Paolo De Coppi

**Author notes:** Correspondence should be addressed to: Paolo De Coppi, MD, PhD NIHR Professor of Paediatric Surgery, Nuffield Professor of Paediatric Surgery, Head of Stem Cells & Regenerative Medicine Section, Developmental Biology& Cancer Programme, UCL Great Ormond Street Institute of Child Health, 30 Guilford Street, London WC1N 1EH, UK. Jan A Deprest, MD, PhD, Fetal Medicine Unit, Department of Obstetrics and Gynaecology, University Hospitals Leuven, Katholieke Universiteit Leuven, B-3000 Leuven, Belgium. Marko Z. Nikolic, MD, PhD, UCL Respiratory, Division of Medicine, University College London, 5 University Street, London WC1E 6JF, UK. These authors contributed equally to this work (Stavros P. Loukogeorgakis, Federica Michielin, Noura Al-Juffali, Julio Jimenez, Soichi Shibuya, Jessica Allen-Hyttinen).

## Abstract

**Rationale:** Congenital diaphragmatic hernia (CDH) results in lung hypoplasia. In severe cases, tracheal occlusion (TO) can be offered to promote lung growth. However the benefit is limited, and novel treatments are required to supplement TO. Vascular endothelial growth factor (VEGF) is downregulated in animal models of CDH and could be a therapeutic target, but its role in human CDH is not known.

**Objectives:** To investigate whether VEGF supplementation could be a suitable treatment for CDH-associated lung pathology.

**Methods:** Fetal lungs from CDH patients were used to determine pulmonary morphology and VEGF expression. A novel human *ex vivo* model of fetal lung compression recapitulating CDH features was developed and used to determine the effect of exogenous VEGF supplementation (Figure 1A). A nanoparticle-based approach for intra-pulmonary delivery of VEGF was developed by conjugating it on functionalized nanodiamonds (ND-VEGF) and was tested in experimental CDH *in vivo*.

**Measurements and Main Results:** VEGF expression was downregulated in distal pulmonary epithelium of human CDH fetuses in conjunction with attenuated cell proliferation. The compression model resulted in impaired branching morphogenesis similar to CDH and downregulation of VEGF expression in conjunction with reduced proliferation of terminal bud epithelial progenitors; these could be reversed by exogenous supplementation of VEGF. Prenatal delivery of VEGF with the ND-VEGF platform in CDH fetal rats resulted in lung growth and pulmonary arterial remodelling that was complementary to that achieved by TO alone with appearances comparable to healthy controls.

**Conclusions:** This innovative approach could have a significant impact on the treatment of CDH.

## Introduction

Pulmonary hypoplasia is a major cause of neonatal mortality (rates of 70-95% perinatally). Congenital diaphragmatic hernia (CDH) is one of the major causes; it is characterised by herniation of abdominal viscera into the thorax resulting in impaired lung development, reduced airway branching/alveolar surface area, and pulmonary hypertension^1,2^. Severe/extreme CDH is associated with mortality in excess of 75%^3^, and can be diagnosed antenatally enabling prenatal therapy. Currently, tracheal occlusion (TO), involving percutaneous balloon insertion into the fetal trachea^4,5^, is the only *in utero* intervention available to fetuses with compromised lung development^6^. Even with TO, mortality remains around 50%^7^, necessitating novel therapies complementary to TO^8^.

CDH animal models have shown impairment of the VEGF pathway. VEGF levels are reduced in hypoplastic lungs from rats with nitrofen-induced CDH^9,10,11,12^, whilst prenatal pharmacological disruption of VEGF signalling resulted in hypoplasia and pulmonary hypertension in rodents and sheep^13,14,15^. Moreover, VEGF administration *in vitro* accelerated growth in nitrofen-induced hypoplastic rat lung explants^16^ and normalised function of pulmonary artery endothelial cells derived from fetal lambs with CDH^17^. However, the role of the VEGF pathway in human CDH pathogenesis has not been investigated. A significant contribution to the CDH pulmonary phenotype arises from the mechanical compression of the herniated organs into the chest cavity. However, in the Nitrofen model, lung development defects occur earlier than herniation^8^ so teratogenic *versus* mechanical effects are difficult to distinguish, whereas surgical models of CDH are limited to late stages of development therefore not accurately recapitulating the pathogenesis^18^.

Although the therapeutic effects of prenatal VEGF administration have been demonstrated in animal models of prematurity-associated respiratory distress syndrome^19,20^, there is currently no *in vivo* evidence that VEGF administration has an effect on CDH. Successful use of exogenous VEGF in this context would necessitate a delivery method that allows VEGF release in a sustained and targeted manner. The latter is particularly important due to the rapid plasma clearance of VEGF, as well as its key role in tumour growth^21^. Nanoparticle-based delivery platforms are currently being investigated in various fields of medicine^22,23,24^, and may also have significant translational potential in the setting of CDH-associated pulmonary pathology. Nanodiamonds (ND; 2-8nm carbon nanoparticles) are one such platform, integrating several properties that are prerequisites for clinical translation allowing conjugation and sustained release of a broad range of therapeutic agents^24,25^.

## Methods

### Mouse and human fetal lung tissue

Whole lungs from E12.5 mouse embryos were obtained as previously reported^26^. Whole fetal lung samples aged 7 – 8 pcw were obtained under permission from NHS Research Ethical Committee (REC reference 18/LO/0822 and 18/NE/0290) and the Joint MRC/Wellcome Trust Human Developmental Biology Resource (www.hdbr.org).

### *3D* printing

For three-dimensional (3D) printing, 10µL of 20% PEG-HCC in PBS were added on Matrigel-embedded lung samples. 3D-printing was performed as previously reported^27^. Confocal microscope settings for crosslinking: pixel resolution 1640×1640, pixel dwell time 0.98μs, zoom 0.9x, laser wavelength 800nm, stack depth 2μm, laser power 80%, 4x immersion objective. Control samples were incubated with the same amount of PEG-HCC gel but without cross linking.

### Production of Nanodiamond-VEGF Conjugates

VEGF conjugation on nanodiamonds (ND) was carried out using a sequential process summarised in Supplementary Methods, Supplementary Results and Supplementary Figure 11A.

### Animals for in vivo study

All experimental protocols were approved by the Ethics committee for Animal Experimentation of the Faculty of Medicine, KU Leuven, Leuven, Belgium (P102/2015) and follow the ARRIVE 2.0 reporting guidelines. Gestational day (E) 19-21 fetuses from time-dated pregnant Wistar rats were used in experiments.

### Generation of CDH

In order to generate fetuses with CDH and hypoplasia, pregnant rats received 100mg of nitrofen (by gavage in 1mL olive oil) on gestational day 9 (E9) as previously described^10,11,28^.

### Sample Size Calculation and Statistical analysis

Results are expressed as mean ± SDEV of biological replicates as indicated in figures. Continuous variables were compared by Mann-Whitney test or One-way ANOVA with Bonferroni post-hoc tests (GraphPad Prism 7.0; San Diego, California, USA). For *in vivo* experiments, sample size calculation was done based on previous studies where TO improved several outcomes by 40% in CDH fetuses^29,30^. We expected an additional effect of 60% over the effect of TO, and thus outcome within the range of healthy fetuses. Using as variables 8 groups, alpha 0.05 and beta 0.05, we calculated a minimum of 8 animals per group.

## Results

### Lung hypoplasia in human CDH fetal lungs is associated with reduced VEGF expression and epithelial proliferation

We found noticeable hypoplasia as early as 18 pcw in post-mortem human CDH tissue compared to healthy lung tissue at matching developmental stages (Figure 1B-C)^31^. We also found significant downregulation of VEGF and its receptor 2 (KDR) in the distal epithelium compared to healthy controls with associated reduction in proliferation (Figure 1D-G). Overall, these data are similar to previous findings using the CDH nitrofen model^9^. We therefore hypothesised that the exogenous supplementation of VEGF to the developing lung could promote distal epithelial proliferation and attenuate lung hypoplasia in CDH fetuses.

**Figure 1.**
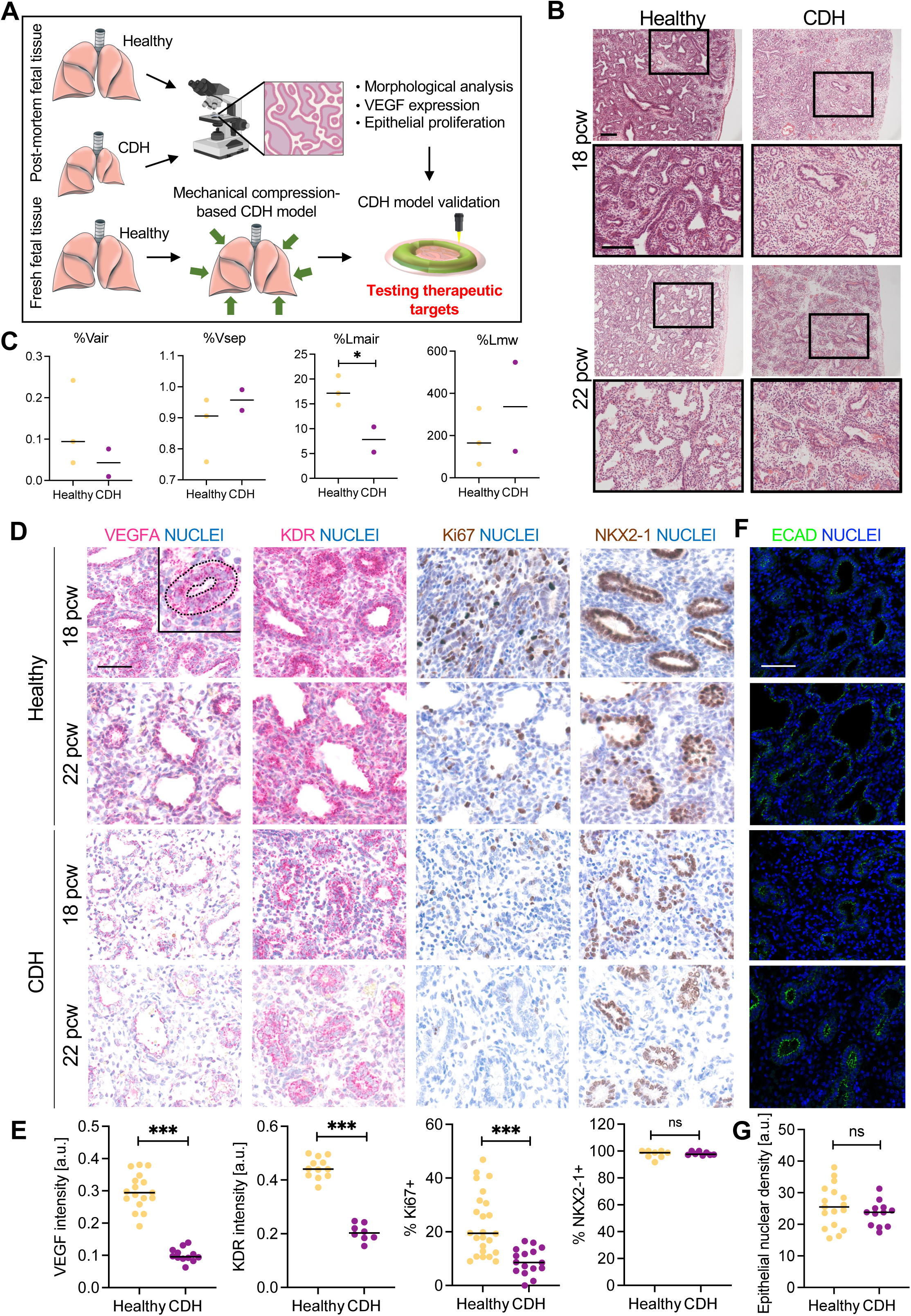
(A) Schematic of the study design based on the analysis of the early-stage CDH fetal lung sections to assess early hypoplasia and VEGF impairment. In parallel, a human compression-based CDH model will be developed to validate the CDH phenotype and test a therapeutical approach to rescue lung hypoplasia. (B) H&E analysis of 18 and 22 pcw post-mortem CDH human lung tissue reveals impaired epithelial development compared to control healthy tissue at matching developmental stages. Scale bar 100μm. (C) Morphometric analysis of CDH and healthy lung tissue at 18-22 pcw reveals decreased values of volume densities of air space (Vair%) and corresponding higher values of volume densities of alveolar septa (Vsep%) in CDH tissues compared to healthy tissue at the same developmental stages. A similar trend can be observed for the mean linear intercept of air space (Lma%) and its corresponding mean linear intercept of septal thickness (Lmw%). (D) Histologic analysis of VEGFA, KDR, Ki67 and NKX2-1 markers in normal and CDH fetal lung tissue at 18 and 22 pcw. Scale bar 100μm. (E) Quantification of epithelial VEGFA and KDR expression reported in D shows a significant decrease for both markers in CDH compared to healthy samples. Correspondingly, the percentage of Ki67-positive epithelial cells is significantly lower in CDH compared to healthy samples, with no significant differences in the number of NKX2-1-positive cells. ***<p=0.001. (F) Immunofluorescence analysis of ECAD in normal and CDH fetal lung tissue at 18 and 22 pcw. Scale bar 100μm. (G) Quantification of epithelial nuclear density based on results in Figure F shows no significant differences between CDH and healthy tissue.

### Development of a human model of fetal lung mechanical compression relevant to CDH

To test this hypothesis, we sought to develop an *ex vivo* model that recapitulates the mechanically constricted environment of the human fetal lung at the earliest stages of CDH. We first exposed embryonic mouse lungs to physical constraints created through hydrogels (PEG-HCC) with well-defined elastic modulus, by leveraging an intravital bioprinting approach^32^, that enables an accurate control over the PEG-HCC crosslinking around the lung with predetermined mechanical properties (Figure 2A). Atomic Force Microscopy (AFM) analysis confirmed the increase in stiffness of the crosslinked PEG-HCC gel compared to Matrigel (Figure 2B). No cytotoxic effects as a direct consequence of the crosslinking have been observed^32,27^. The formation of a mechanically printed barrier successfully confined the expansion of mouse lung tissue compared to non-compressed controls (Figure 2C; Supplementary Video 1). The area occupied by the left lobe was significantly reduced and branching morphogenesis was impaired in the confined lungs compared to control. In addition, H&E staining revealed morphological alterations of the compressed lungs with noticeable reduced airway space (Figure 2D-E). We then analysed cell proliferation within the pulmonary epithelial compartment and found that the fraction of ECAD/EdU double positive cells in compressed lungs was reduced (Figure 2F). In addition, bulk RNA-sequencing analysis demonstrated altered transcriptional activity in the compressed lungs compared to controls, including VEGF-A downregulation (Figure 2G; Supplementary Figure 1). DEGs network analysis showed enrichment of GO-BP categories related to inflammatory and immune responses^33,34^ (Figure 2H; Supplementary Spreadsheet 1). To better understand how the compressed lung is subjected to the mechanical load as a consequence of the confinement, we developed an *in-silico* analysis based on the finite element method (FEM). We assumed a distinct proliferation rate between distal and proximal areas of the tissue, that was validated by the experimental observations of surface growth (Supplementary Figure 2A-C). Hydrostatic stress maps showed an overall increase of the mechanical load in the compressed lung with a noticeable difference between proximal and distal areas, suggesting bud tips are the most subjected to compression (Supplementary Figure 2D).

**Figure 2.**
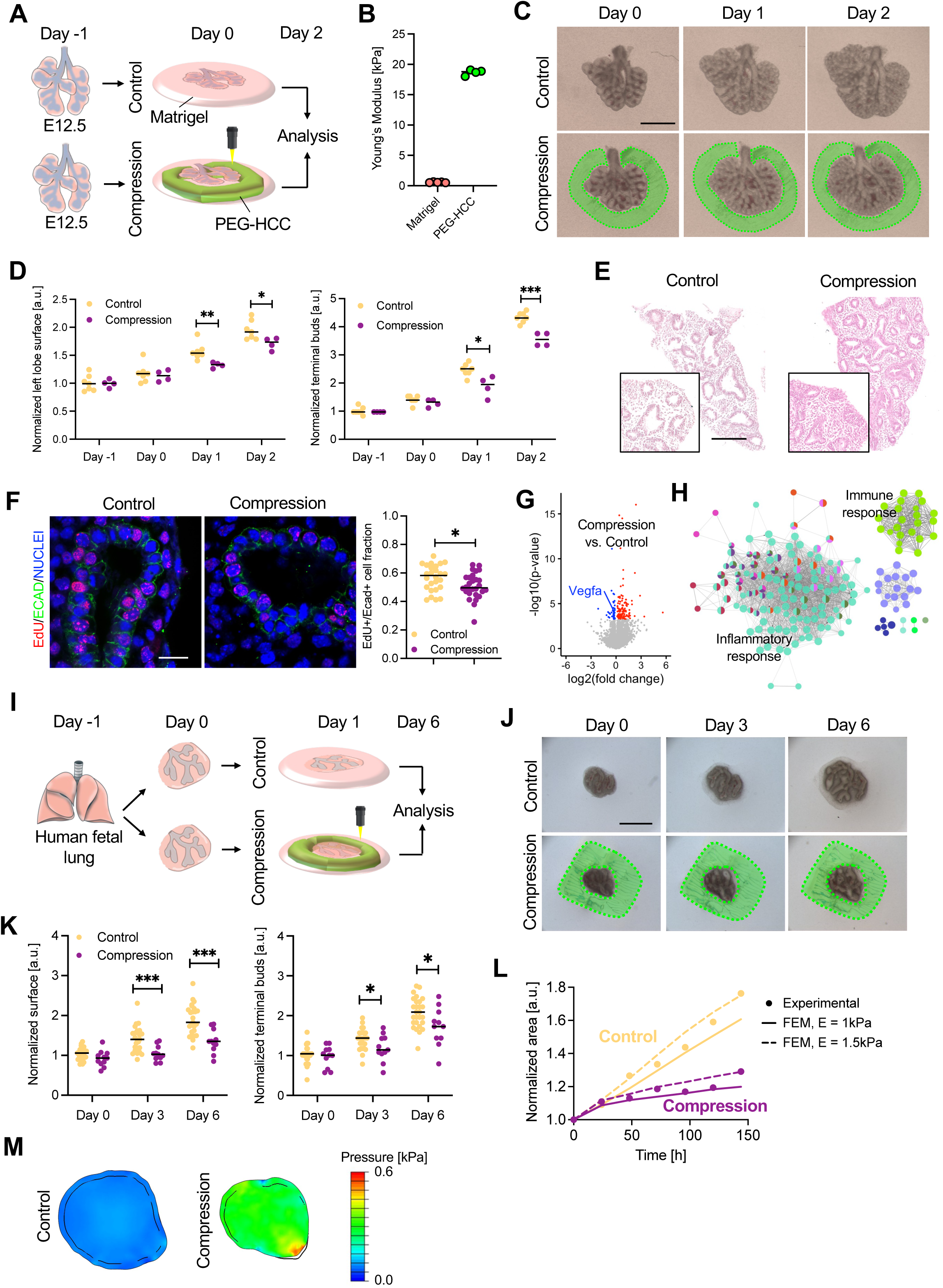
(A) Experimental setup of the *ex vivo* model of mouse fetal lung compression during pseudoglandular stage. Freshly-isolated mouse fetal lungs (E12.5) were embedded in Matrigel drops and cultured in Air-liquid Interphase (ALI). The next day (Day 0), a 20% Cumarin HCC-conjugated 8-arm-polyethylene glycol gel (PEG-HCC) solution was dispersed into the Matrigel drop and selectively crosslinked under a two-photon microscope to surround the lungs and inhibit their growth over a total of 2 days. Control lungs had the same amount of PEG-HCC solution added to the Matrigel, but this was not crosslinked with the two-photon laser. (B) Atomic force microscopy analysis confirms the stiffness of the gel in culture conditions after two weeks from printing is approximately 20-fold higher compared to Matrigel. (C) Time-lapse imaging of lung growth under physical confinement (Compression) compared to control. Scale bar 100μm. (D) Quantification of left lobe surface area (left) and terminal buds (right) in control and compressed lungs after two days of mechanical confinement shows significant impairment of lung growth and branching in compressed lungs. *N*=6 embryos. *<p=0.05; **<p=0.01; ***<p=0.001. (E) H&E staining reveals qualitatively noticeable alterations of the fetal tissue morphology in control and compressed lungs after two days of mechanical confinement. *N*=2 embryos. Scale bar 100μm. (F) EdU incorporation assay reveals a decreased proliferation of distal epithelial cells in the compressed lungs compared to control. *N*=3 embryos *<p=0.05. (G) Volcano plot of bulk RNA-sequencing analysis of compressed vs control lungs after two days of mechanical confinement. Differentially expressed genes (DEGs) have been identified with FC>1.2 and FDR<0.05. DEGs analysis shows downregulation of *Vegfa* in compressed lungs. (H) Gene network analysis of overexpressed genes in the compressed lungs reveals upregulation of categories related to inflammatory and immune response. (I) Experimental setup of the *ex vivo* model of human fetal lung compression during pseudoglandular stage. The compression is achieved by the physical confinement of fetal lung distal tissue through the 3D-printing of the PEG-HCC gel after 24 hours of culture in Matrigel. The tissue is then cultured *ex vivo* for seven days in the presence or absence of physical confinement. (J) Representative time-lapse imaging of human lung tissue growth under physical confinement (Compression) compared to control. *N*=4 donors. Scale bar 100μm. (K) Quantification of human lung tissue surface area (left) and terminal buds (right) in control and compressed lungs after six days of mechanical confinement. *N*=4 donors. *<p=0.05; **<p=0.01; ***<p=0.001. (L) FEM analysis validation based on the time-course experiment in Supplementary Figure 2E. Comparison of normalized area of the lung sample over time for experimental data and numerical values from FEM analysis. The area is normalized to the geometric configuration at Day 0. FEM analysis was developed adopting two values of the Young’s modulus to account for the uncertainty about the stiffness of the human lung tissue. (M) Pressure field prediction. Representative pressure field (kPa) for control and compressed lung fragments at Day 6, assuming a Young’s modulus of 1.5 kPa for the human lung tissue.

We then translated the developed *ex vivo* CDH model for human tissue investigation, by using early pseudoglandular stage human fetal lung fragments (Figure 2I; Supplementary Methods). We first verified tissue growth and viability and observed branching morphogenesis and spontaneous peristalsis up to 2 weeks in culture (Supplementary Video 2). To mimic mechanical compression, we used PEG-HCC as before to 3D-print a mechanical barrier surrounding the lung (Figure 2I). At day 7 we observed a significant reduction in the surface area of compressed lung fragments, as well as reduced branching morphogenesis (Figure 2J,K). FEM analysis adapted to the human model (Supplementary Figure 2E,F) showed a good representation of experimental data assuming a Young’s modulus comprised between 1 and 1.5 kPa (Figure 2L). Hydrostatic stress maps confirmed an approximate 3-fold increase of the mechanical load in the compressed tissue compared to control (Figure 2M). Overall, these data suggest that the mechanical confinement at an early stage is sufficient to mimic some key features of CDH pathogenesis, mainly affecting distal epithelial proliferation.

### VEGF expression and cell proliferation are impaired in compressed bud tip progenitor cells and can be rescued by exogenous supplementation of VEGF in the human fetal lung compression model

To further investigate the effects of mechanical compression on the human fetal lung tissue, and on the bud tip cells in particular, we conducted a single-cell RNA-sequencing (scRNA-seq) analysis of compressed and control lung fragments from 3 donors. We captured and identified over 30,000 cells across 3 biological replicates, including all expected cell types with no significant differences in both compressed and control lungs^35^ (Figure 3A; Supplementary Figure 3A-D). Within the epithelial compartment, we identified expected distal epithelial cells (comprised of tip, stalk, and airway progenitors), as well as neuroendocrine, and very few peripheral nervous system (PNS) cells. We also found a sizeable cluster of undifferentiated cells adjacent to the distal epithelial fraction which have not been described in previous single-cell studies using fresh tissue; these strongly expressed stretch-associated genes^36,37^ in addition to distal epithelial markers, and were not associated with the presence of the mechanical constraint. We therefore identified these as ‘stretched epithelial’ cells and hypothesise that their presence is due to an artifact of *ex vivo* culture, as swelling of distal tips in culture was occasionally observed (Supplementary Figure 4). We focused our investigation on the *SOX9*/*TESC* double-positive tip cells^35^. GO-BP analysis revealed several pathways significantly upregulated in compressed tips (Figure 3B-C); many of these signals are lost when looking at the epithelial compartment as a whole (Supplementary Figure 3E,F), indicating a heightened effect of compression on tip cells as predicted. Furthermore, we saw a reduction in known proliferation signatures^38^ and VEGF-family genes in the presence of mechanical stimuli (Figure 3D).

**Figure 3.**
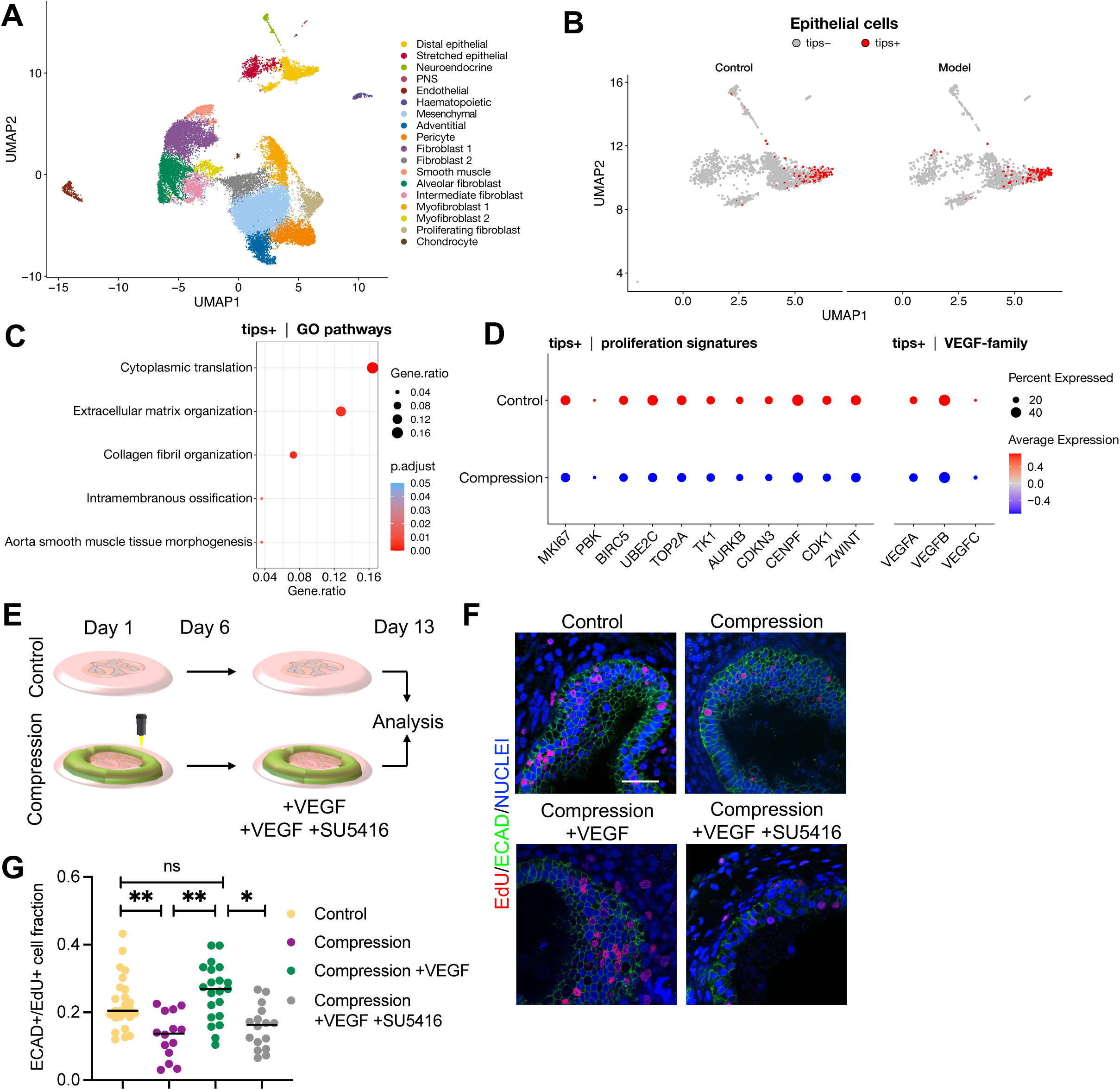
(A) UMAP visualization of 32630 fetal lung cells, n=3 biological samples. Leiden clustering revealed 18 distinct cell types within mesenchymal, epithelial, endothelial, and hematopoietic fractions. (B) Magnified UMAP of the epithelial compartment cells. Distal tip cells, “tips+”, were identified by co-expression of known tip markers (*SOX9*+, *TESC*+); the remaining cells comprise non-tip epithelium, “tips-”. (C) Gene Ontology enrichment analysis reveals differential expression of several gene profiles in tips+ cells of the CDH model. (D) Relative expression of 11 proliferation-associated genes and 3 VEGF-family genes between control and CDH model tips. (E) Experimental set up of the human *ex vivo* model to test exogenous VEGF supplementation. After 7 days of culture in basal medium, compressed human lung fragments were treated with recombinant VEGF for an additional 7 days, using the specific KDR/Flk1 (VEGF receptor 2) inhibitor SU5416 as negative control. (F) Representative pictures of EdU proliferation assay after 7 days of VEGF treatment (Compression +VEGF) compared to Control, Compression only and compressed samples treated with the VEGF-specific inhibitor SU5416. Immunostaining for E-CADHERIN (E-CAD) was used to identify epithelial cells. Scale bar 100μm. (G) Quantification of the number of ECAD/EdU double-positive cells after 7 days of VEGF treatment shows a significant increase in proliferation in VEGF-treated samples (Compression +VEGF) compared to not treated samples (Compression). Proliferation in samples after VEGF treatment is also significantly higher than proliferation in samples treated with the VEGF inhibitor SU5416. *N*=2 donors. *<p=0.05; **<p=0.01; ***<p=0.001.

We leveraged the human model to test whether the noted reduction in proliferative markers in the distal tips resulted in actual mitotic impairment, and if supplementation of exogenous VEGF, administered at optimal concentrations based on previous studies using murine lungs^16^ (Supplementary Figure 5A-C), could mitigate this effect (Figure 3E). While mechanical confinement significantly impairs distal epithelial proliferation, VEGF supplementation significantly rescued epithelial proliferation to levels comparable to control (uncompressed) lungs; the same effect was not obtained in the presence of SU5416, confirming the specificity of the proposed treatment (Figure 3F,G). Overall these data suggest VEGF could be a target for restoring epithelial proliferation in the distal lung when this is exposed to mechanical confinement.

### Exogenous VEGF supplementation promotes lung growth and ameliorates airway abnormalities in the nitrofen rat model of CDH

To validate our hypothesis and test the therapeutic value of VEGF *in vivo*, we then sought to determine whether exogenous VEGF supplementation would be salutary (over and above TO) against pulmonary hypoplasia in the nitrofen rat model of CDH. We developed a nanoparticle (nanodiamond; ND) VEGF delivery platform (ND-VEGF) to allow spatially-and temporally-targeted, gradual release of VEGF in fetal lungs *in vivo*. The effect of ND-VEGF in this setting was compared to that of TO alone, as well as VEGF administered to the fetal lung of TO animals as a solution (free VEGF). Finally, we investigated the possible mechanism of action of VEGF in this setting, by co-administering the KDR/Flk1 inhibitor SU5416^39,40^. Experimental groups are summarised in Figure 4A and Supplementary Methods. Ultrasound assessment at E19 was 100% accurate in detecting fetuses with CDH as confirmed at harvesting (E21; Figure 4B). Excluding animals in the healthy control group, only fetuses with ultrasound-detected CDH at E19 were included in the study (CDH incidence: 40%). There was no difference in fetal survival between experimental groups (Supplementary Figure 6). Fetal body weight at E21 was significantly reduced in sham-treated CDH animals compared to healthy controls and was not affected by any of the interventions (data not shown). Biodistribution studies identified ND-VEGF only within the distal airways of CDH rats at E21, with evidence of retention of VEGF by E-Cadherin-positive epithelial cells (Figure 4C). Histopathological analysis of fetal and maternal tissues demonstrated no toxic effects of empty ND, free VEGF, and ND-conjugated VEGF. In addition to lack of bronchial oedema and other pulmonary side-effects, *in utero* intratracheal administration of free and ND-conjugated VEGF did not result in microscopic abnormalities or neovascular growth in the placenta, as well as fetal or maternal organs (Supplementary Figure 7,8).

**Figure 4.**
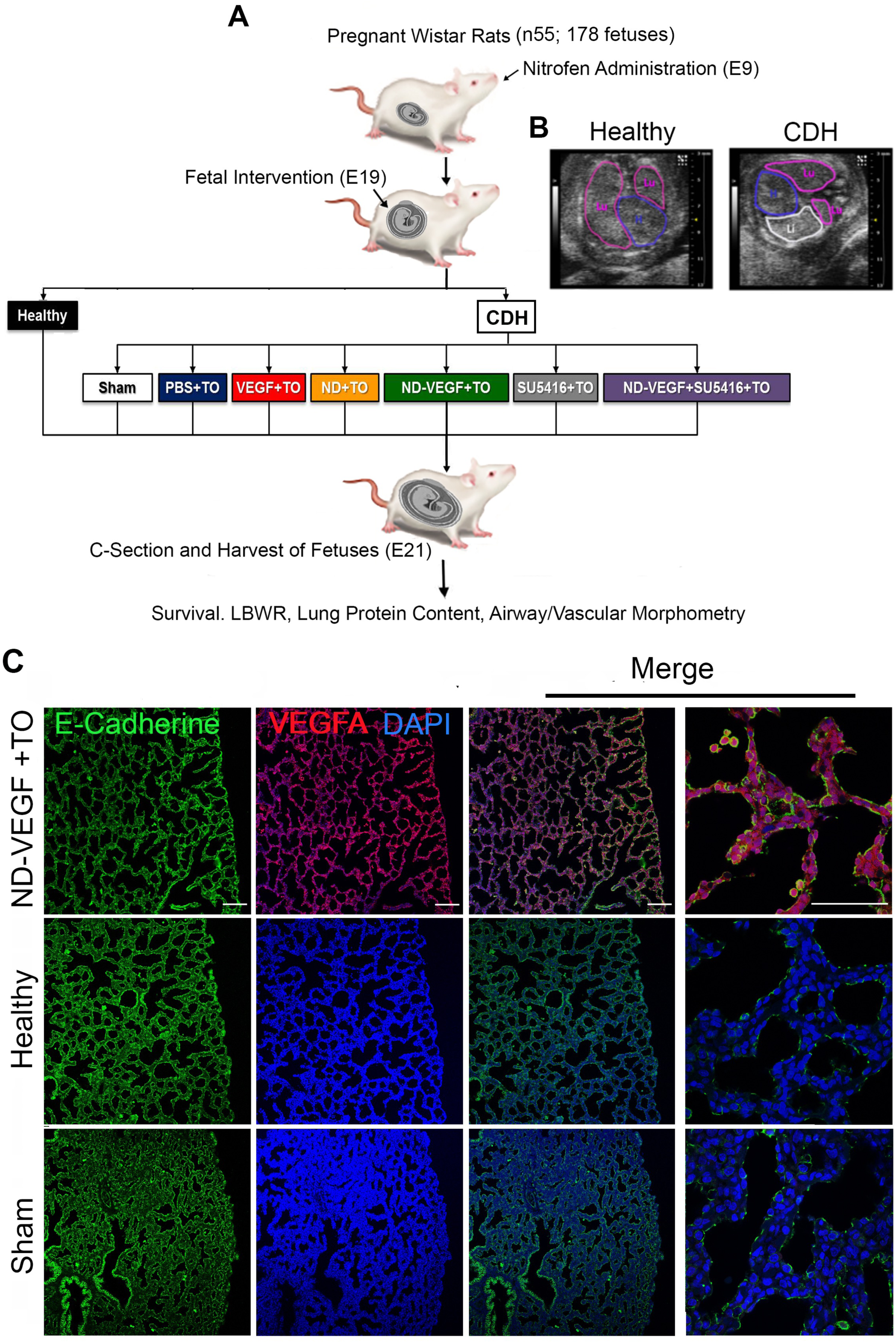
(A) Experimental design of in vivo intervention studies in fetal rats using VEGF-loaded nanodiamonds (ND-VEGF). There were eight experimental groups: healthy control (non-congenital diaphragmatic hernia/CDH; olive oil gavage-fed mothers), sham intervention (surgery but no injection or tracheal occlusion/TO in CDH fetuses from nitrofen-fed mothers), PBS+TO (intra-tracheal vehicle injection at E19 followed by TO in CDH fetuses), free VEGF+TO (free VEGF injection followed by TO), ND+TO (unconjugated ND-NH2 injection followed by TO), ND-VEGF+TO (ND-VEGF injection followed by TO), SU5416+TO (KDR/Flk1 inhibitor SU5416 injection followed by TO) and ND-VEGF+SU5416+TO (ND-VEGF and SU5416 co-injection followed by TO). Animals in the groups with ND-VEGF administration received 13.4±1.9μg ND and 100ng VEGF, while animals in ND+TO and VEGF+TO received the same amounts of ND-NH2 and free thiolated-VEGF in 50μL of PBS. The concentration of the KDR/Flk1 inhibitor SU5416 in the injection solution used in SU5416+TO and ND-VEGF+SU5416+TO groups was 120μg/mL. (B) Confirmation of the presence of diaphragmatic defect in E18 fetal rats using micro ultrasound (Lu/pink outline: lung; H/purple outline: heart; Li/white outline: liver). (C) Representative confocal microscopy image demonstrating localisation of ND-VEGF in airways at E21, following E19 intra-tracheal administration with TO, with evidence of uptake/retention of VEGF by E-Cadherin expressing alveolar epithelial cells (E-Cadherin: green; VEGF: red; DAPI: blue; magnification columns 1-3: x20; magnification column 4: x63).

There were obvious macroscopic differences in lungs harvested from animals in different experimental groups. Lungs in the sham group were the smallest, those in the ND-VEGF+TO group were the biggest, while the size of lungs in other groups was in-between that observed in sham and ND-VEGF+TO animals (Figure 5A). Lung-to-body weight ratio (ratio of fetal lung to total body weight; LBWR) and total lung protein measurements content in untreated CDH fetuses were markedly reduced compared to healthy controls. TO resulted in a significant increase in both LBWR and protein content compared to sham (Figure 5B,C). *In utero* intra-tracheal administration of ND-VEGF followed by TO had effects that were complementary to those of TO alone, and led to a lung size comparable to that observed in healthy controls (Figure 5B,C). This therapeutic effect of ND-VEGF was not observed when free VEGF or empty ND were administered, and was abrogated by co-administration of the KDR/Flk1 inhibitor SU5416. In agreement with previously published data, nitrofen-induced CDH resulted in marked abnormalities in airway morphology that were partly improved by TO (Figure 5D,E; Supplementary Figure 9). TO resulted in partial improvement in these parameters (Figure 5E; Supplementary Figure 9). ND-VEGF had salutary effects on lung morphology over and above to those achieved by TO alone and normalised distal airway maturation in CDH animals (Figure 5E), but there was no concomitant normalisation of the number of alveolar saccules (Supplementary Figure 9). Similar to what we observed in terms of lung growth, the therapeutic benefit of ND-VEGF was not observed when free VEGF or empty ND were administered and was abrogated by co-administration of SU5416 (Figure 5E). SPC expression in ND-VEGF+TO animals was similar to healthy controls and significantly higher than in sham-treated animals (Figure 5G). SU5416 co-administration inhibited ND-VEGF-induced alveolar epithelial cell maturation and TO alone (or in conjunction with free VEGF or empty ND) did not affect SPC expression, which was found to be similar to that observed in the sham group (Figure 5G).

**Figure 5.**
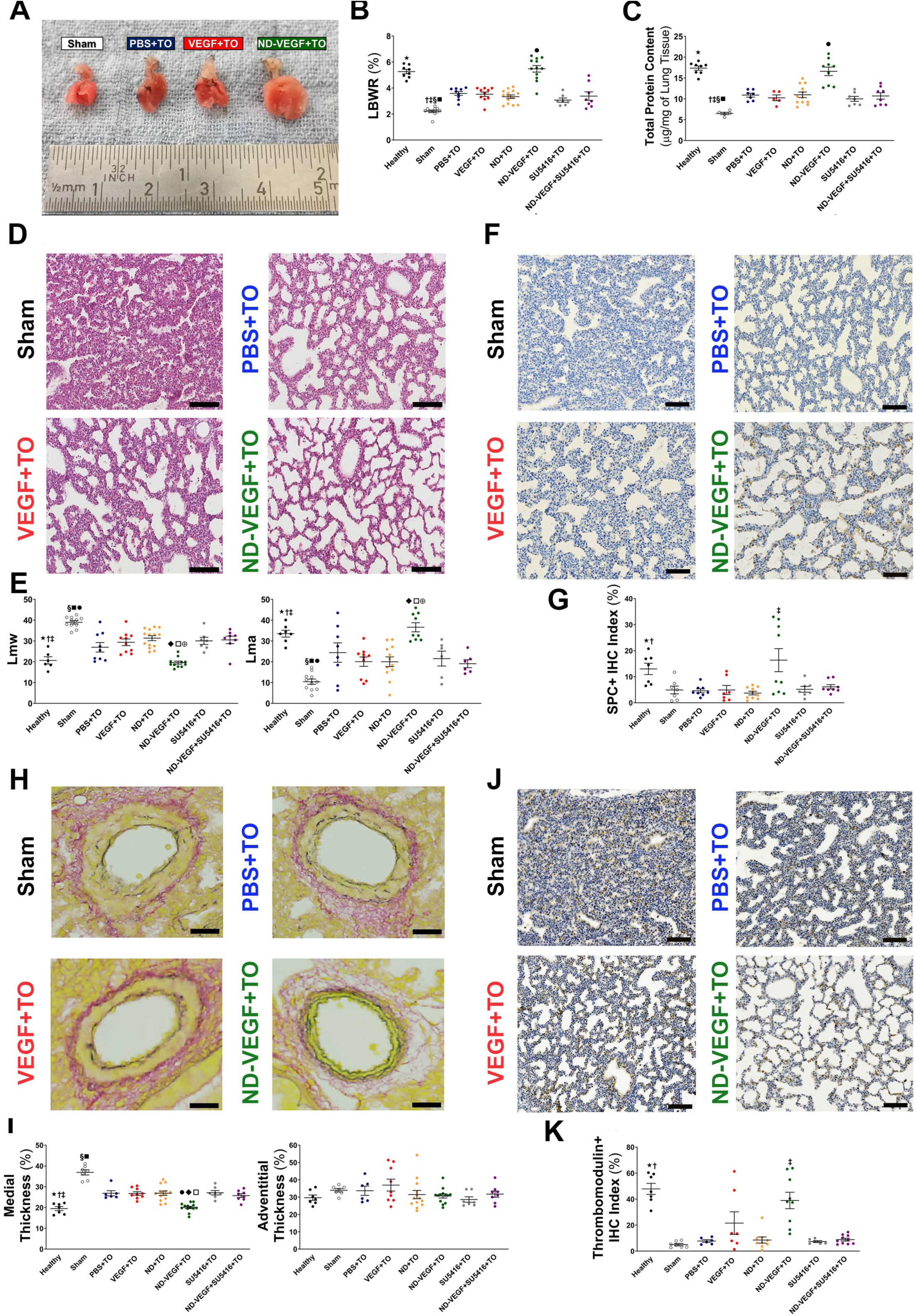
(A) Representative images of E21 lungs from rats in the intervention study (left to right: sham, PBD+TO, VEGF+TO, ND-VEGF+TO). (B) Summary of lung-to-body weight ratio data (expressed as a percentage; p<0.0001 vs Sham, PBS+TO, ND+TO, VEGF+TO, SU5416+TO and ND-VEGF+SU5416+TO; † p<0.0001 vs. PBS+TO & ND-VEGF+TO; ‡ p<0.001 vs. ND+TO & VEGF+TO; § p<0.01 vs. ND-VEGF+SU5416+TO; ■p<0.05 vs SU5426+TO; ●p<0.0001 vs. PBS+TO, ND+TO, VEGF+TO, SU5426+TO & ND-VEGF+SU5416+TO). (C) Summary of total protein content data (expressed as µg/mg of lung tissue; ★p<0.0001 vs Sham, PBS+TO, ND+TO, VEGF+TO, SU5416+TO and ND-VEGF+SU5416+TO; † p<0.0001 vs. PBS+TO & ND-VEGF+TO; ‡ p<0.001 vs. ND+TO & VEGF+TO; § p<0.01 vs. ND-VEGF+SU5416+TO; ■p<0.05 vs SU5426+TO; ●p<0.0001 vs. PBS+TO, ND+TO, VEGF+TO, SU5426+TO & ND-VEGF+SU5416+TO). (D) Representative images of sections of E21 rat lungs from animals included in the in vivo intervention studies. Tissues were stained with hematoxylin and eosin (H&E) and were used for subsequent airway morphometry analyses. (E) Summary of mean linear intercept of wall transection length (Lmw) data (★p<0.0001 vs. Sham; † p<0.01 vs. VEGF+TO, ND+TO, SU5426+TO and ND-VEGF+SU5426+TO; ‡ p<0.05 vs. PBS+TO; § p<0.0001 vs. ND-VEGF+TO; ■p<0.001 vs. PBS+TO; ●p<0.01 vs. VEGF+TO, ND+TO, SU5416+TO and ND-VEGF+SU5416+TO; ♦p<0.001 vs. ND+TO, SU5416+TO and ND-VEGF+SU5416+TO; □p<0.01 vs. VEGF+TO; ⨁p<0.05 vs. PBS+TO), and mean linear intercept of parenchymal airspace (Lma) data (★p<0.0001 vs. Sham; † p<0.01 vs. VEGF+TO, ND+TO, SU5426+TO and ND-VEGF+SU5426+TO; ‡ p<0.05 vs. PBS+TO; § p<0.0001 vs. ND-VEGF+TO; ■ p<0.001 vs. PBS+TO; ●p<0.01 vs. VEGF+TO, ND+TO, SU5416+TO and ND-VEGF+SU5416+TO; ●p<0.001 vs. ND+TO, SU5416+TO and ND-VEGF+SU5416+TO; ●p<0.01 vs. VEGF+TO; ⴲp<0.05 vs. PBS+TO). (F) Representative images of lung sections from in vivo intervention studies stained for surfactant protein C (SPC; brown) using immunohistochemistry (IHC). (G) Summary of SPC+ IHC index (ratio of the number of SPC+ cells over total cell number in pre-determined areas of lung parenchyma; expressed as percentage) data (★p<0.0001 vs. Sham, PBS+TO, ND+TO, SU5416+TO and ND-VEGF+SU5416+TO; † p<0.01 vs. VEGF+TO; ‡ p<0.0001 vs. Sham, PBS+TO, VEGF+TO, ND+TO, SU5416+TO and ND-VEGF+SU5416+TO). (H) Representative images of sections of E21 rat lungs from animals included in the in vivo intervention studies. Miller’s elastic staining was performed, and stained sections were used for subsequent vascular morphometry analyses (peripheral arterioles 30-50µm diameter). (I) Summary of medial thickness data (expressed as a percentage of the external diameter of the blood vessel) (★p<0.0001 vs. Sham; **†** p<0.01 vs. VEGF+TO, ND+TO, SU5426+TO and PBS+TO; **‡** p<0.05 vs. ND-VEGF+SU5426+TO; **§** p<0.0001 vs. ND-VEGF+TO; ◆p<0.001 vs. PBS+TO, VEGF+TO, ND+TO, SU5416+TO and ND-VEGF+SU5416+TO; ●p<0.01 vs. ND+TO; ♦p<0.001 vs. PBS+TO and ND-VEGF+SU5416+TO; □ p<0.01 vs. ND-VEGF+SU5426+TO), and summary of adventitial thickness data (expressed as a percentage of the external diameter of the blood vessel). (J) Representative images of lung sections from in vivo intervention studies stained for thrombomodulin (brown) using immunohistochemistry (IHC). (K) Summary of thrombomodulin+ IHC index (ratio of the number of thrombomodulin+ cells over total cell number in pre-determined areas of lung parenchyma; expressed as percentage) data (★p<0.0001 vs. Sham, PBS+TO, ND+TO, SU5416+TO and ND-VEGF+SU5416+TO; **†** p<0.01 vs. VEGF+TO; **‡** p<0.001 vs. Sham, PBS+TO, ND+TO, SU5416+TO and ND-VEGF+SU5416+TO).

### Exogenous VEGF supplementation ameliorates pulmonary vascular abnormalities in the nitrofen rat model of CDH

Nitrofen-induced CDH was associated with increased muscularization of peripheral pulmonary arterioles (diameter: 30-50µm), as well as reduced overall alveolar vascularization (Figure 5H,J). Medial thickness (MT) in the sham group was higher (Figure 5H,I) and thrombomodulin expression in alveolar tissue was lower (Figure 5J,K) compared to healthy controls (Figure 5I,K). TO alone attenuated pulmonary arteriolar muscularization to levels that were still significantly higher than in healthy controls (Figure 5I), but had no effect on alveolar vascularization (Figure 5K). Similar to what we observed for lung growth and morphology, ND-VEGF in conjunction with TO was the only intervention that resulted in near-normalisation of both arterial muscularization (Figure 5H,I) and overall alveolar vascularization (Figure 5J,K), but these effects were reversed when SU5416 was co-administered (Figure 5K,I). Pulmonary arteriolar adventitial thickness was similar in all experimental groups (Supplementary Figure 10).

## Discussion

Herein we demonstrate that exogenous VEGF supplementation, coupled with TO has the potential to rescue lung hypoplasia and pulmonary vascular abnormalities in CDH. As failure of diaphragm closure likely starts manifesting around 6 post-conception weeks^8^, we collected post-mortem CDH tissue at the earliest available fetal stages and confirmed impaired VEGF expression in the distal epithelium of CDH lungs. We also found altered lung morphometry reflecting impaired branching morphogenesis, along with reduced epithelial proliferation. We developed an *ex vivo* model using human fetal lung tissue and confirmed that mechanical compression alone is sufficient to induce morphological and transcriptional signs of lung hypoplasia in the absence of chemical or genetic manipulation. In contrast to previous studies using lung organoids to model CDH loads^41^, here we propose a whole fetal lung tissue compression model including all cell types present at a stage at which CDH likely starts manifesting *in vivo*. A preliminary characterization of the model using embryonic mouse lung evidenced impaired branching morphogenesis and reduced epithelial proliferation, confirming results in human CDH tissue. In addition, we found an overall transcriptional upregulation of inflammatory and immune response genes, along with downregulation of *Vegf*. Interestingly, an inflammatory response has been detected both in the Nitrofen model^33^ and in human cord blood from CDH fetuses^42^. We developed an FEM-based analysis and quantified mechanical stress, to confirm that distal areas are stiffened compared to control. We then leveraged the model to investigate compression of human tissue, using distal fragments isolated from fetal lungs and showing for the first time that mechanical compression alone decreases proliferation in bud tip progenitors. Single-cell RNA-sequencing revealed a reduced expression of *VEGF* in compressed epithelial bud tip progenitors, possibly reflecting the significant VEGF impairment found later in CDH lungs. Although the mechanism through which mechanical alterations lead to VEGF impairment is unknown, we speculate that an autocrine/paracrine VEGF signalling, previously described in epithelial development and in different types of cancer^43,44,45^ could be involved.

Although the therapeutic effects of prenatal VEGF administration have been demonstrated in prematurity-associated respiratory distress syndrome models^19,20^, there is currently no *in vivo* evidence for an effect of VEGF administration in CDH. Such administration would require tight temporal and spatial control to emulate normal development. Nanodiamonds integrate several properties that make them suitable in this setting, including innate biocompatibility, scalability, precise particle distribution, a high surface-area-to-volume ratio, a near-spherical aspect ratio and an easily adaptable carbon surface for bio-agent attachment^46^. Their use as a biocompatible drug delivery platform is effective in pre-clinical treatment of chemo-resistant tumours thanks to prolonged and sustained drug delivery *in situ*^25^. In our innovative system, VEGF covalently bound to a nanodiamond carrier allowed sustained release within the fetal lung. As expected, fluorescently labelled nanoparticles showed a non-uniform distribution with a granular aspect in the cytoplasm of pulmonary epithelial cells suggesting cellular uptake by endocytosis, in contrast to small molecules that often diffuse through the cell membrane^47^. The containment of the nanoparticles within endosomes prevents ejection by the cells’ efflux pump mechanism, but also inhibits penetration deep into the cytosol; this is likely an advantage as VEGF functions primarily extracellularly in fetal airway development^48^. Importantly, even epithelial cells exposed to increasing concentrations of ND for prolonged periods showed no changes in metabolic activity, proliferation, or apoptosis. Moreover, there were no visible changes to cell adhesion or cytoskeletal structure assessed using fluorescence imaging, and we could not detect any cytotoxicity induced by ND.

We tested a therapeutic formulation where VEGF was coupled to ND structures. Chemical modification of the VEGF protein into a sulfhydryl form did affect biological activity in the well-established CAM assay^49^. We also investigated the biological effects of VEGF in the Nitrofen rat model^50^. Pulmonary VEGF expression is reduced in CDH^10^ but upregulated after TO, corresponding to substantial lung growth^29^. We confirmed this, and further demonstrated that the addition of injected substances did not increase embryonic rat mortality. Injection of unmodified VEGF did not further supplement the effect of TO, but injection of VEFG-ND markedly improved LBWR and protein/lung weight ratio to control levels. This effect is completely abolished when a specific VEGF inhibitor^39,40^ is injected alongside the VEGF-ND complex, as also confirmed by lung morphometry. As hypothesized, gradual release of VEGF by ND-VEGF affected lung vasculature with normalization of overall pulmonary parenchymal vascularisation (assessed by expression of the endothelial marker thrombomodulin)^51^, as well as medial thickness of peripheral pulmonary arteries. TO alone only partially induces these changes, and the effect disappears when a VEGF inhibitor is added, suggesting that the beneficial effects of supplemental VEGF in reversing lung hypoplasia require sustained release. We believe that this finding goes beyond CDH treatment and could be also adopted in other perinatal lung diseases in which VEGF plays a key role^20,52^.

There are limitations of the study. Firstly, the mechanism of action is unclear and while specific, it is uncertain whether VEGF acts on pulmonary epithelial, endothelial cells or both. Furthermore, we did not measure ND-VEGF nanoparticle leakage to other organs, although we did not see any leakage in non-pulmonary tissues in our biodistribution study. Pulmonary tropism could be enhanced by modifying the carrier to bind to cell membranes of organ-specific cells^53^. Finally, local lung toxicity remains unclear *in vivo*, however it seems not to be significant in higher species^54^. This should not be underestimated because nanoparticles are taken up by local immune cells which may play critical roles in the induction and regulation of pulmonary immunity or inflammation^47^.

In conclusion, we have demonstrated the role of VEGF in maturation of human lung epithelium and its impairment in CDH. Both *in vitro* and *in vivo* data show that sustained VEGF rescued the affected lungs. Delivery of growth factors by nanocarriers could have a place in perinatal lung therapy with potential scalability for clinical use, but this will need to be tested in a larger animal model for CDH and TO.

## Author Contributions

S.P.L., F.M.: conception and design, financial support, provision of study material, collection of data, data analysis/interpretation, manuscript writing, final approval of manuscript; N.A.J., J.J., S.S: conception and design, collection of data, data analysis/interpretation, manuscript writing, final approval of manuscript; J.A-H: collection of data, data analysis/interpretation, manuscript writing, final approval of manuscript; P.E., A.S.N.A: data analysis/interpretation, final approval of manuscript; J.D.: manuscript writing, final approval of manuscript; E.N., P.M.: collection of data, data analysis/interpretation, final approval of manuscript; A.T., S.K., A.K..: data analysis/interpretation, manuscript writing, final approval of manuscript; M.P.: provision of study material, final approval of manuscript; C.F.: data analysis/interpretation, manuscript writing, final approval of manuscript; A.D.: data analysis/interpretation, manuscript writing, final approval of manuscript; N.K., D.S., T.X.: collection of data, final approval of manuscript; N.S.: provision of study material, final approval of manuscript; M.G.: analysis/interpretation, final approval of manuscript; S.E.: data analysis/interpretation, final approval of manuscript; J.T.: data analysis/interpretation, manuscript writing, final approval of manuscript; C.L.: data analysis/interpretation, final approval of manuscript; P.P.: data analysis/interpretation, manuscript writing, final approval of manuscript; P.C.: financial support, provision of study material, data analysis/interpretation, manuscript writing, final approval of manuscript; F.R.: data analysis/interpretation, final approval of manuscript; S.J.: financial support, provision of study material, data analysis/interpretation, manuscript writing, final approval of manuscript; M.Z.N., N.E.: conception and design, financial support, provision of study material, data analysis/interpretation, manuscript writing, final approval of manuscript; J.D.: conception and design, financial support, provision of study material, data analysis/interpretation, manuscript writing, final approval of manuscript, P.D.C.: conception and design, financial support, provision of study material, data analysis/interpretation, manuscript writing, final approval of manuscript.

## Supporting information

Supplementary methods & results

## Acknowledgements

This work and S.P.L. was supported by a Wellcome Trust post-doctoral fellowship for MB/PhD graduates (ref. number: 098539/Z/12/Z). F.M. was supported by a NIHR BRC Catalyst Fellowship and a Rosetrees Trust grant (Seedcorn2020/100052). P.D.C. and F.M. are supported by Building Respiratory Epithelium and Tissue for Health (BREATH) Consortium for Lung Regeneration (LongFond, Dutch). S.S. was supported by Japan society for the promotion of science overseas research fellowships (310072). This work and J.J. was supported by the European Commission via its Erasmus Joint Doctoral program (2013-0040; this publication represents the views of the authors and the European Commission cannot be held responsible from any use which may be made from the information contained therein). P.D.C. is supported by the National Institute for Health Research and the Wellcome Trust/Engineering and Physical Sciences Research Council “Guided Instrumentation for Fetal Therapy and Surgery” programme. J.D. is a Clinical Researcher of the Flanders Research Foundation (FWO Vlaanderen; 8012.07) and is also partly funded by GOSH Children’s Charity. The research on CDH at the KU Leuven is funded by the KU Leuven (C32/17/054), CDH-UK, the Wellcome Trust (WT101957) and the Engineering and Physical Sciences Research Council (EPSRC) (NS/A000027/1). M.Z.N. acknowledges funding from a MRC Clinician Scientist Fellowship (MR/W00111X/1), Rosetrees Trust (M899) and from the Rutherford Fund Fellowship allocated by the MRC UK Regenerative Medicine Platform 2 (MR/5005579/1). J.A-H. acknowledges funding from the University College LondonBirkbeck MRC Doctoral Training Programme.

