## Supplementary Figures for "Prenatal VEGF Nano-Delivery Reverses Congenital Diaphragmatic Hernia-Associated Pulmonary Abnormalities"

***Supplementary Figure 1***

Principal component analysis (PCA) of the indicated samples. Each dot represents a single replicate.

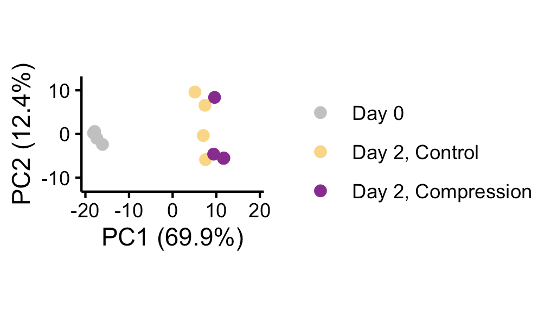

***Supplementary Figure 2***

(A) Time-course of mouse lung growth under physical confinement used for FEM-based model development and validation. Scale bar 500μm. (B) Reconstruction of lung geometries at t = 24 h (equivalent to Day 0 in Figure 2C) based on a different cell proliferation rate between proximal and distal areas as evidenced by EdU incorporation assay. Higher proliferation in the distal area was observed both in ECAD-positive epithelial and ECAD-negative mesenchymal cells. (C) Model validation based on the comparison of normalized area of the lung over time for experimental data and numerical values from FEM analysis. The area is normalized to the configuration at t = 24 h. (D) Prediction of the magnitude displacement field (mm) and pressure field (kPa) at t = 60 h, as estimated by FEM analysis. Low proliferating zone and high proliferating zone are separated by a black solid line. (E) Time-course of human lung fragments growth under physical confinement or not, used for FEM-based model development and validation. Scale bar 500μm. (F) Reconstruction of tissue geometries at Day 0 based on a homogeneous cell proliferation rate.

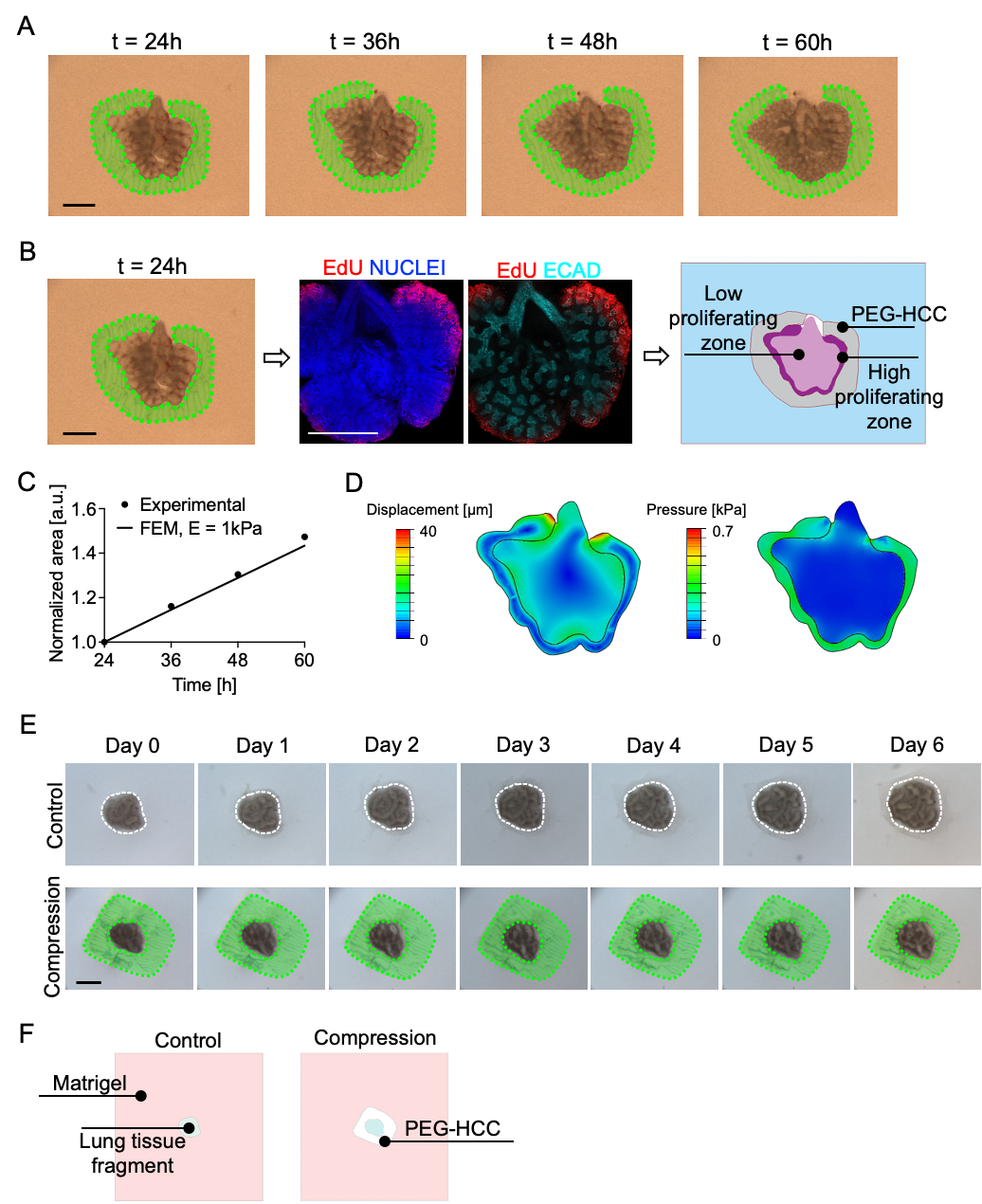

***Supplementary Figure 3***

(A) UMAP of captured cells and (B) annotated cell-type proportions, coloured by replicates. (C) Number of unique transcripts per cell within each annotated cell-type cluster for both control and compressed samples. (D) Proportion of control and compressed cells within the three largest broad cluster annotations (Endothelial, Epithelial, and Mesenchymal). (A-D) No significant deviations in cell type, quantity, or quality in compressed vs control cultured lungs have been observed, although the proportion of each biological and experimental replicate within each of the cell-type clusters was prone to some variation. (E) Pseudo-bulk DE analysis followed by Gene Ontology enrichment shows the biological processes in which differentially expressed genes between control and compressed cells are involved for each broad cluster annotation. (F) Comparison of average gene expression between control and compressed cells for each broad cluster within key top altered pathways.

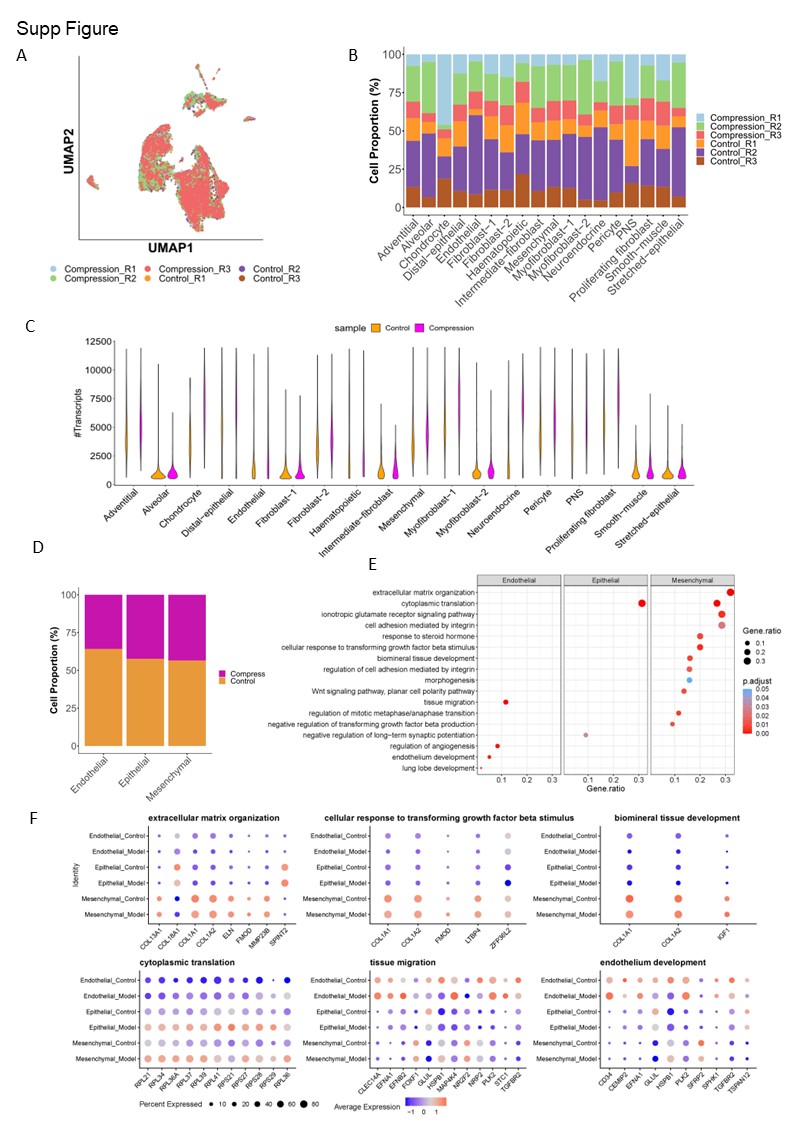

***Supplementary Figure 4***

Example of swelling of distal tips that was occasionally observed in the ex vivo culture of human fetal lung fragments. Scale bar 0.5 mm.

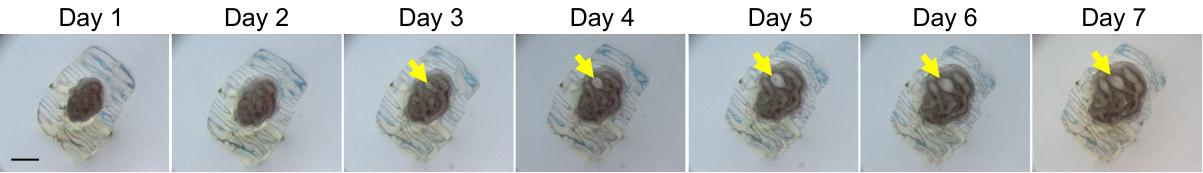

***Supplementary Figure 5***

(A) Experimental set up of VEGF treatment to identify a suitable range of VEGF concentrations that could induce epithelial proliferation. (B) Representative pictures of EdU proliferation assay at Day 13 after treatment with different VEGF concentrations. Immunostaining for E-CADHERIN (E-CAD) was used to identify epithelial cells. Scale bar 100μm. (C) Quantification of the number of ECAD/EdU double-positive cells at Day 13 shows a significant increase in proliferation for a concentration of 50ng/mL. *<p=0.05.

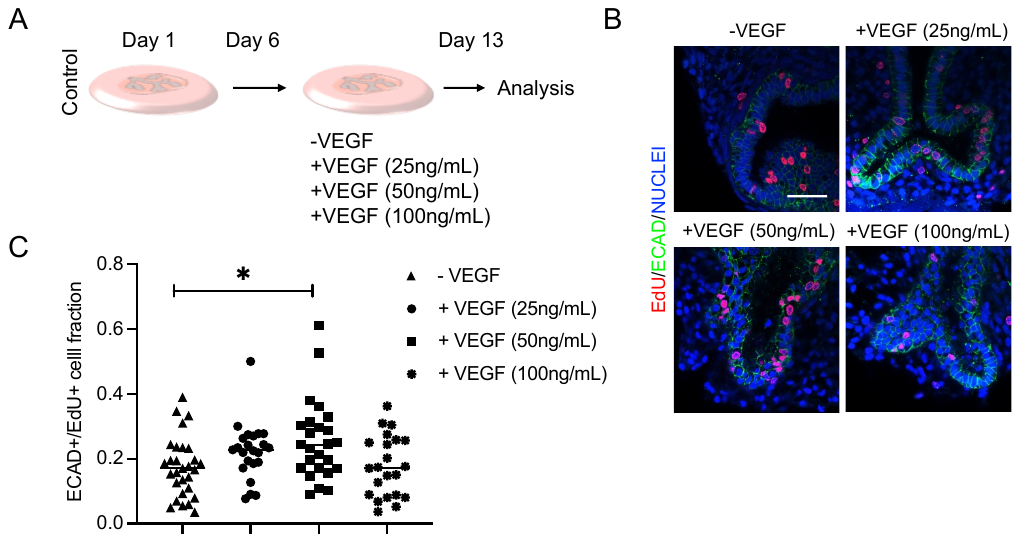

***Supplementary Figure 6***

Survival of fetal rats included in the *in vivo* study (E21; proportion of total injected).

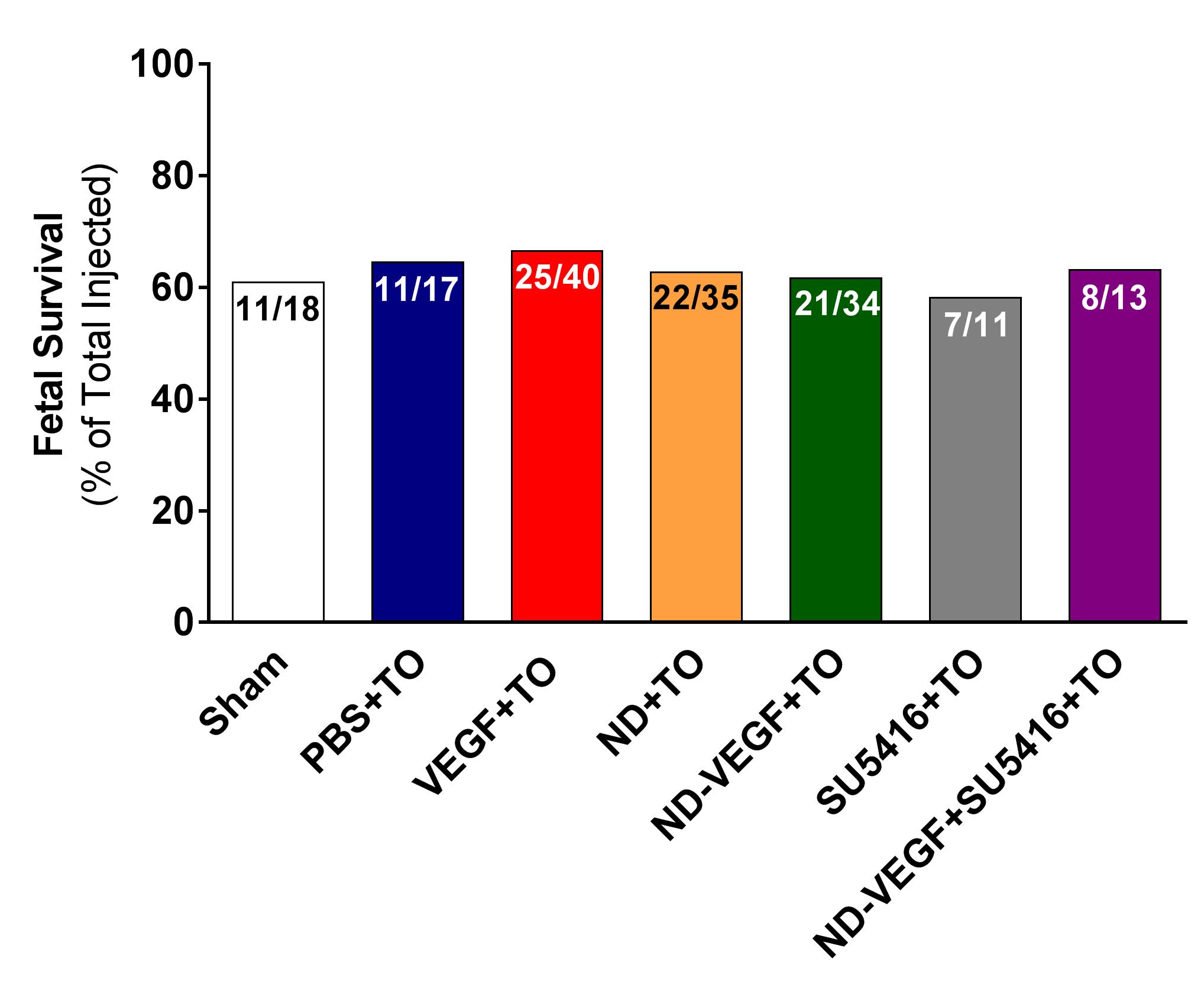

***Supplementary Figure 7***

Representative report from histopathological analysis of tissues from fetus that received in utero administration of vascular endothelial growth factor-loaded nanodiamonds.

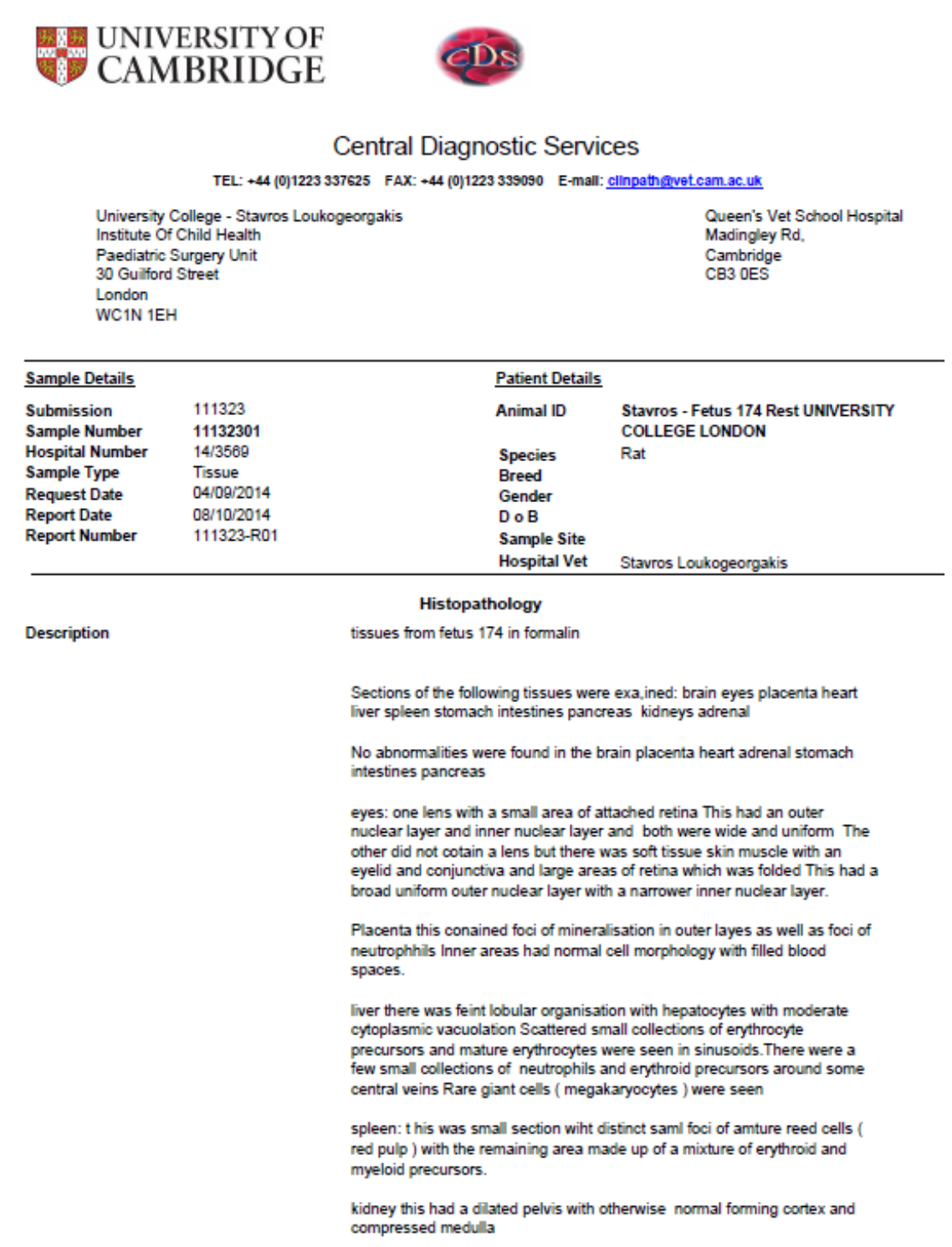

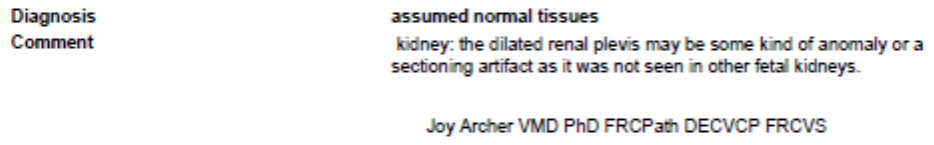

***Supplementary Figure 8***

Representative report from histopathological analysis of tissues from mother the fetuses of which received in utero administration of vascular endothelial growth factor-loaded nanodiamonds.

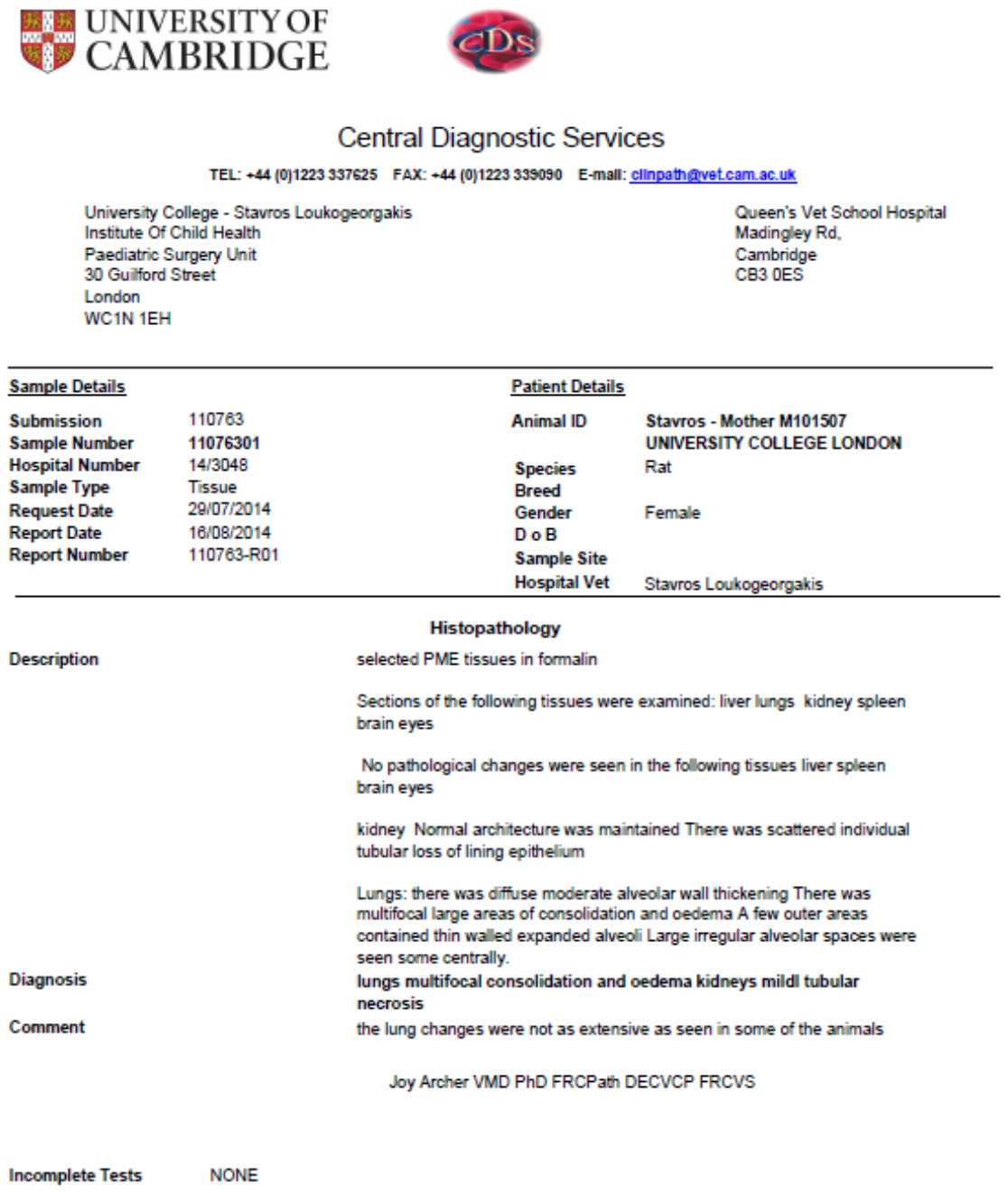

***Supplementary Figure 9***

Summary of mean terminal bronchiolar density (MTBD) data from *in vivo* intervention studies (p<0.0001 vs. Sham; † p<0.001 vs. ND+TO and ND-VEGF+SU5416+TO; ‡ p<0.05 vs. PBS+TO, VEGF+TO and ND-VEGF+TO; § p<0.05 vs. PBS+TO, VEGF+TO, ND+TO, ND-VEGF+TO, SU5416+TO and ND-VEGF+SU5416+TO).

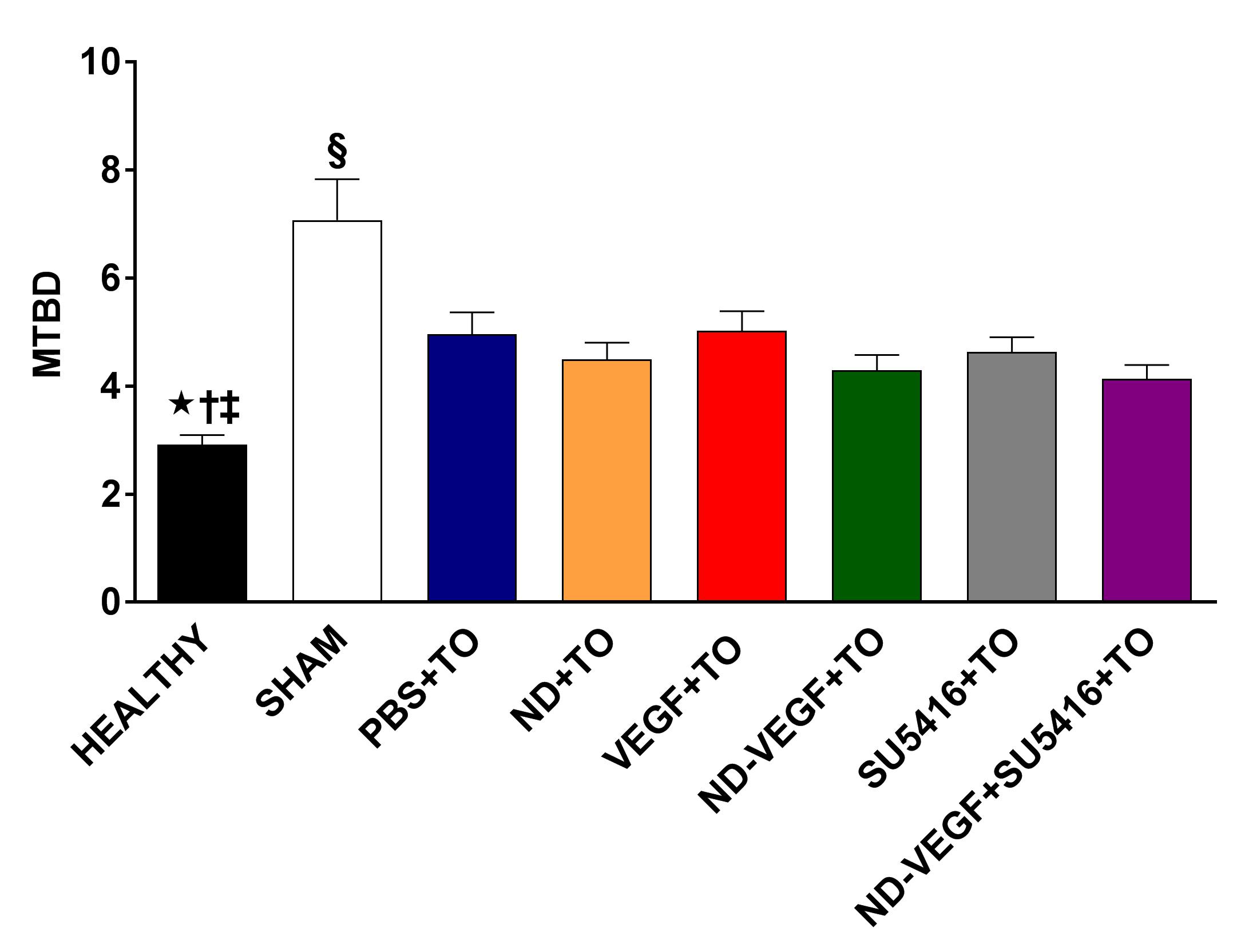

***Supplementary Figure 10***

Summary of adventitial thickness data (expressed as a percentage of the external diameter of the blood vessel) from in vivo intervention studies.

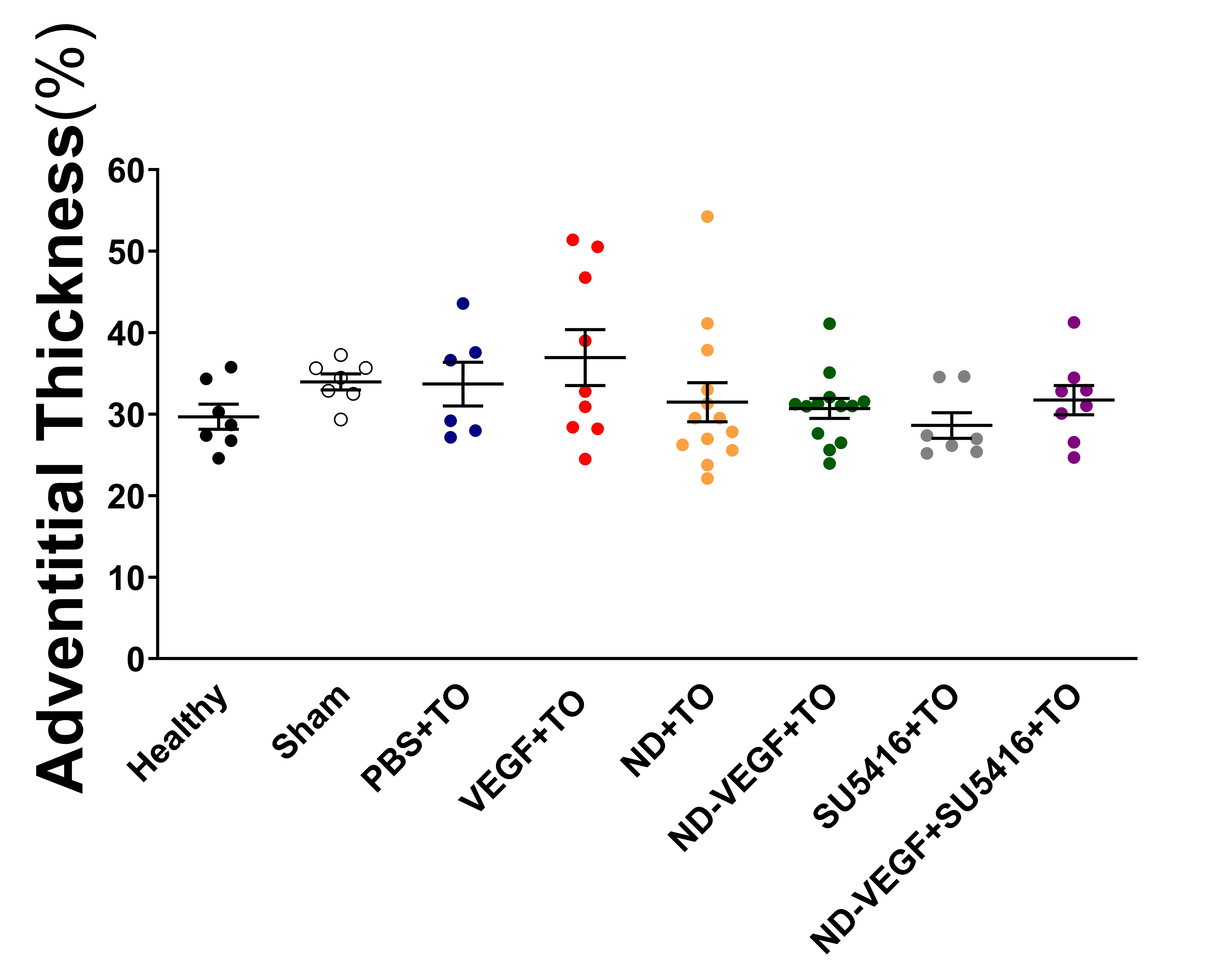

***Supplementary Figure 11***

Nanodiamond characterisation, cellular uptake and toxicity. (A) Scheme for loading of vascular endothelial growth factor (VEGF) on nanodiamonds (ND) (SPDP: Sulfosuccinimidyl 6-(3'-[2-pyridyldithio]-propionamido)hexanoate). (B) Release profile of VEGF from VEGF-loaded nanodiamonds (ND-VEGF). (C) Representative images from chorioallantoic membrane (CAM) vessel proliferation assays comparing the biological activity of stock VEGF (left) to that of thiolated-VEGF used for conjugation to ND (right). (D) Summary results from CAM assays comparing stock and thiolated-VEGF. (E) Transmission electron microscopy (TEM; representative images) of ND with a primary amine group (ND-NH_2_; unconjugated/raw ND; left) and ND-VEGF (right). (F) Flow-cytometric (FACS) analysis of uptake of Alexa Fluor® 488 (AF488; ND-AF488) by human pulmonary epithelial cells. (G) Summary results of EDU (5-ethynyl-2’-deoxyuridine) assays (FACS) to assess effect of addition of increasing concentrations of ND-NH_2_ to culture media on proliferation of human pulmonary epithelial cells. (H) Representative confocal microscopy images demonstrating heterogeneous cytoplasmic distribution of ND-AF488 suggestive of endocytosis (H33258: Hoechst 33258 nuclear stain). (I) Summary results of Alamar Blue assays assessing the effect of addition of increasing concentrations of NF-NH_2_ in the culture media of human pulmonary epithelial cells on their metabolic activity. (J) Representative confocal microscopy images from bromodeoxyuridine / 5-bromo-2'-deoxyuridine (BRDU) assays. (K) Summary results of propidium iodide (PI)/annexin V assays (FACS) assessing the toxicity (early/late apoptosis, necrosis) of co-culture of human pulmonary epithelial cells with ND-NH_2_.

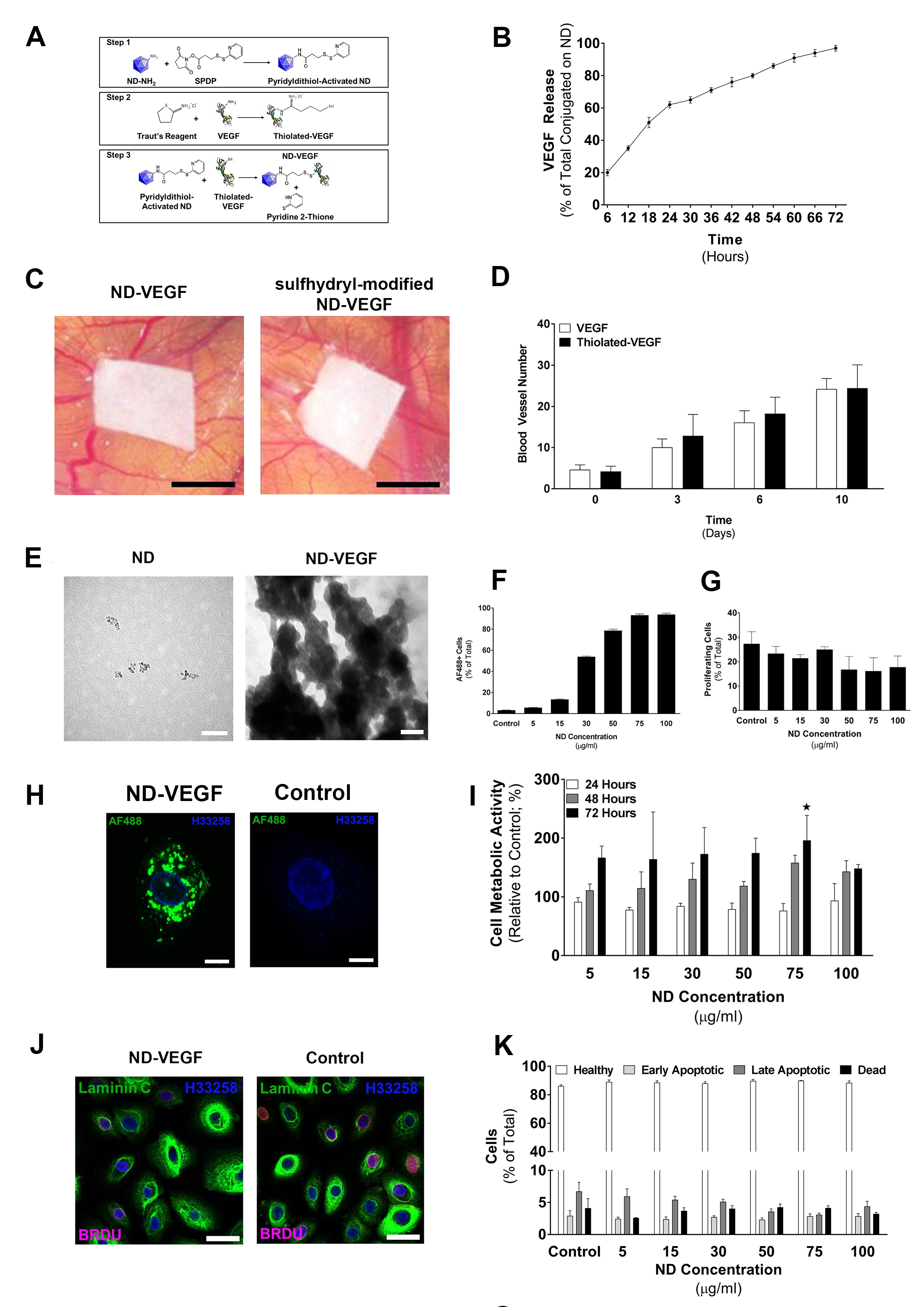

***Supplementary Figure 12***

Synthesis scheme of Alexa Fluor® 488-labelled nanodiamonds. ND: nanodiamond; AF: Alexa Fluor®; NHS: N-hydroxysuccinimide.

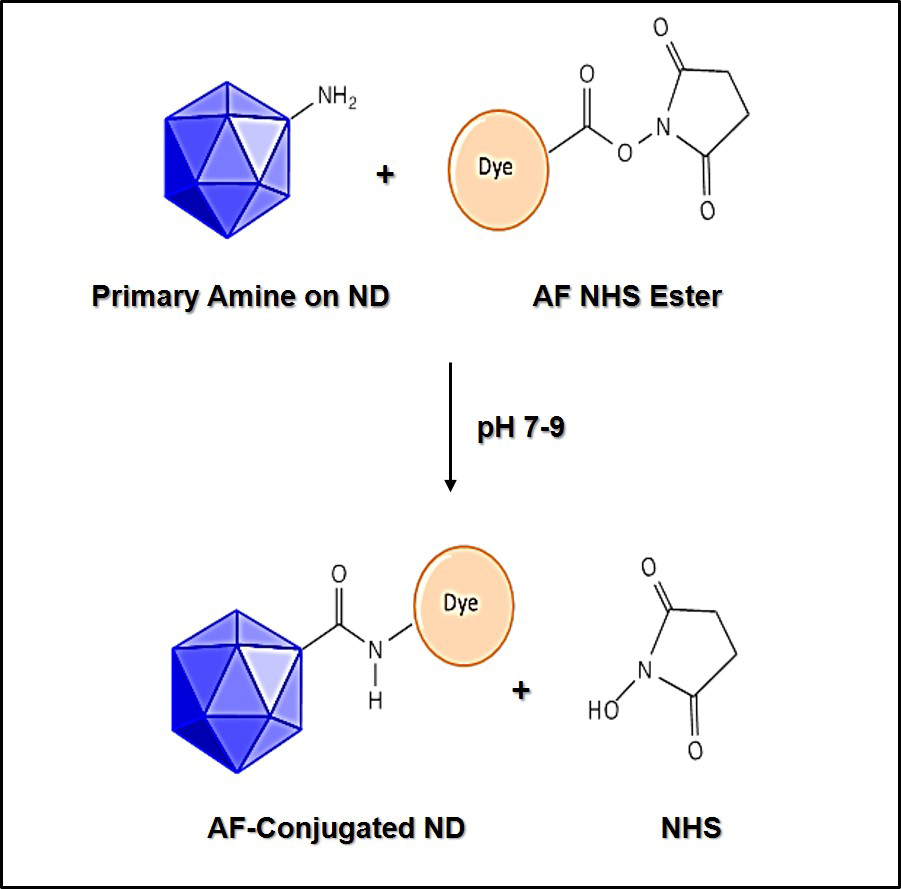

***Supplementary Figure 13***

In vivo toxicity and biodistribution of Alexa Fluor® 488-labelled nanodiamonds following in utero intra-tracheal administration in fetal rats. (A) Experimental design of in vivo toxicity and biodistribution studies using Alexa Fluor® 488-labelled nanodiamonds (ND-AF488). There were four experimental groups: ND-AF488 with tracheal occlusion (TO), ND-AF488 without TO (to determine whether TO is required for uptake of ND in the fetal lung), free AF488 with TO, and free AF488 without TO. Animals in ND-AF488 groups (with/without TO) received 3.9±0.1μg ND and 3μg AF488 (concentration in injection solution of 77.3±1.8μg/ml and 60μg/mL respectively), while animals in AF488 groups (with/without TO) received the same amount of free AF488 in 50μL of PBS. (B) Representative images of surgical procedure for in utero intra-tracheal injection in E19 fetal rats, followed by TO (+: 30G needle inserted in fetal trachea for administration of ND-AF488; *: titanium micro-clip applied to the fetal trachea to achieve TO). (C) Summary of fetal survival data from in vivo toxicity and biodistribution studies. (D) Representative confocal microscopy image demonstrating uptake and retention of ND-488 in airways at E21, following E19 intra-tracheal administration with TO. (E) Representative confocal microscopy image demonstrating minimal uptake and retention of ND-AF488 in airways at E21, following E19 intra-tracheal administration without TO (DAPI: 4′,6-diamidino-2-phenylindole). (F) Representative confocal microscopy image confirming presence of ND-AF488 in distal pulmonary airways, with evidence of uptake by surfactant protein C (SPC) expressing type II alveolar epithelial cells. (G) Representative confocal microscopy image demonstrating lack of retention of AF488 in the lungs of fetuses injected with free dye (in solution; same amount to that carried on ND-AF488).

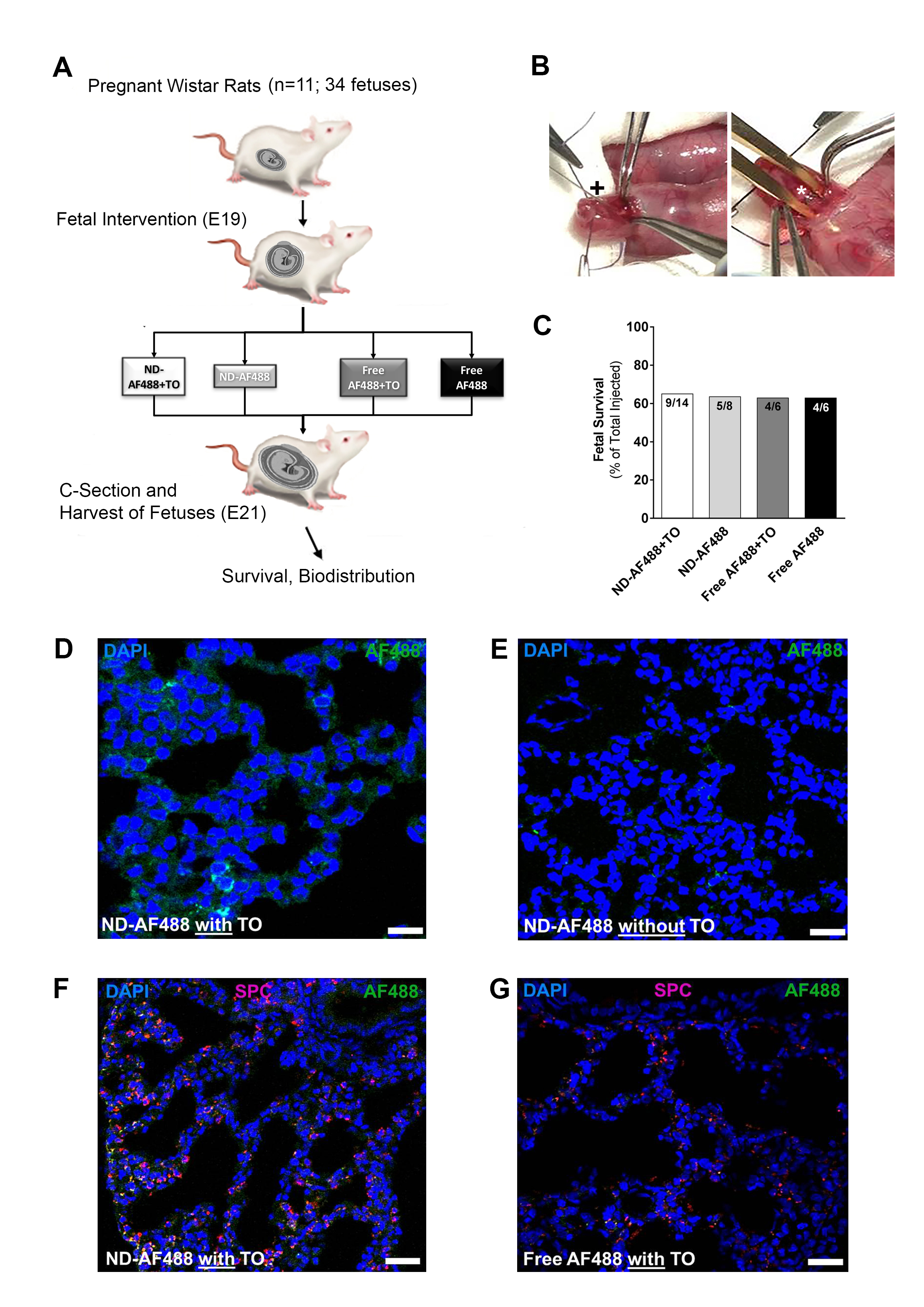

***Supplementary Figure 14***

Confirmation of conjugation of Alexa Fluor® 488 on nanodiamonds by ultraviolet-visible spectroscopy. ND-NH2: Nanodiamonds with primary amine groups; ND-AF488: Alexa Fluor® 488-labelled nanodiamonds.

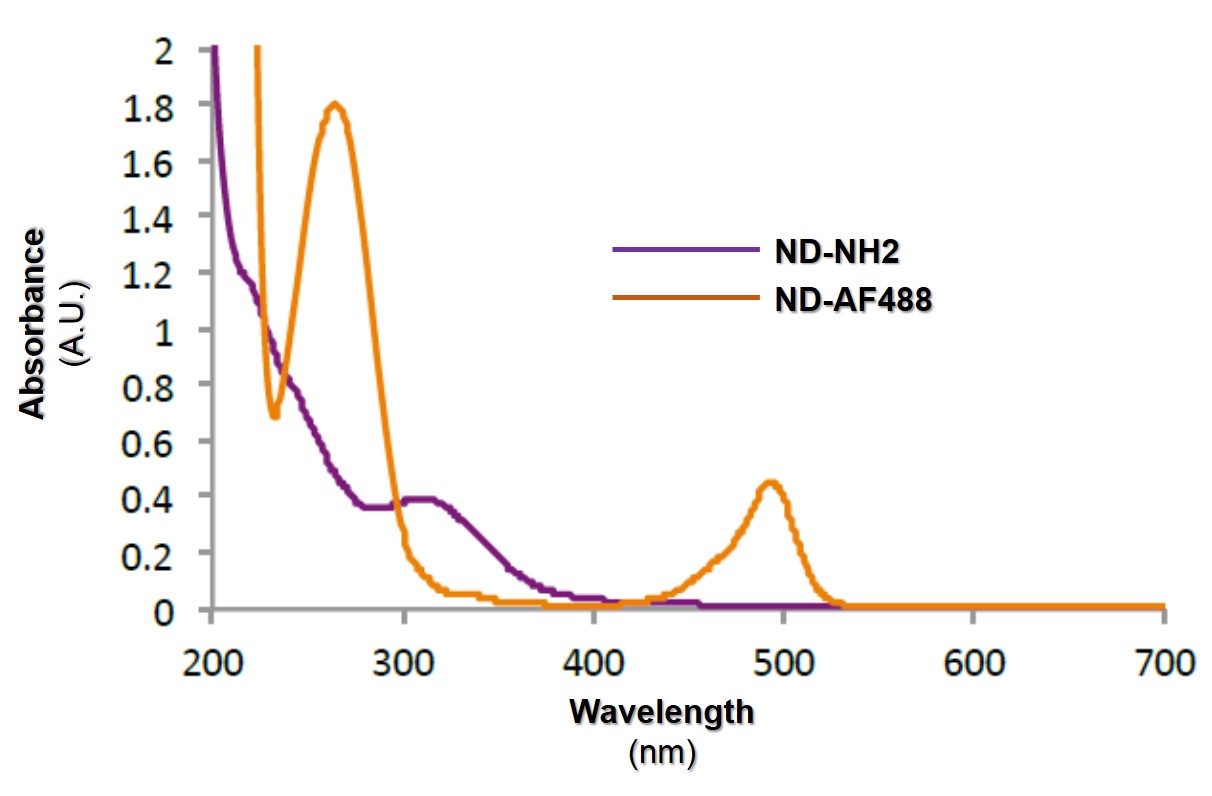

***Supplementary Figure 15***

Confirmation of successful completion of individual steps in the preparation process of vascular endothelial growth factor-loaded nanodiamonds by ultraviolet-visible spectroscopy. ND-COOH; Nanodiamonds with primary carboxyl groups; ND-NH2: Nanodiamonds with primary amine groups; ND-VEGF: VEGF-loaded nanodiamonds.

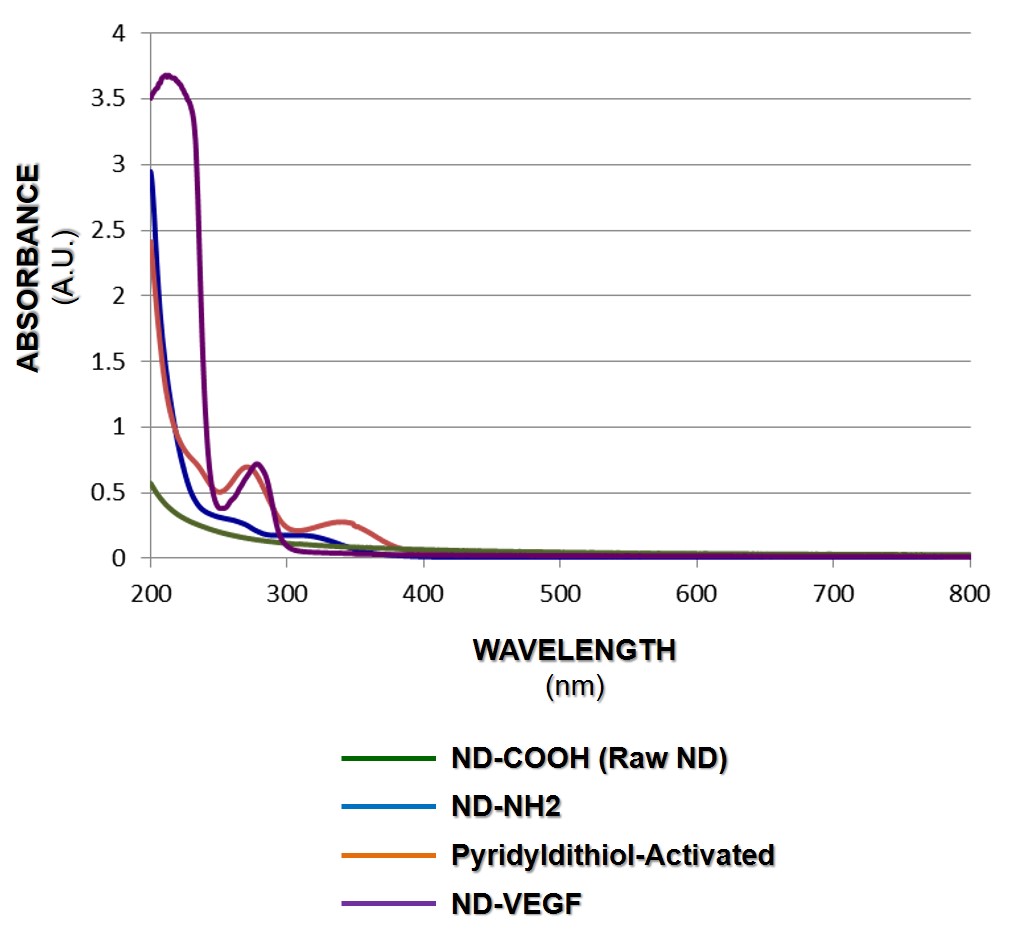

***Supplementary Figure 16***

Confirmation of coupling of sulfhydryl-to-amine cross-linker Sulfosuccinimidyl 6-(3'-[2-pyridyldithio]-propionamido)hexanoate on nanodiamonds by Fourier transform infrared spectroscopy. ND-NH2: Nanodiamonds with primary amine groups; SPDP: Sulfosuccinimidyl 6-(3'-[2-pyridyldithio]-propionamido)hexanoate; ND-SPDP: Pyridyldithiol-activated nanodiamonds

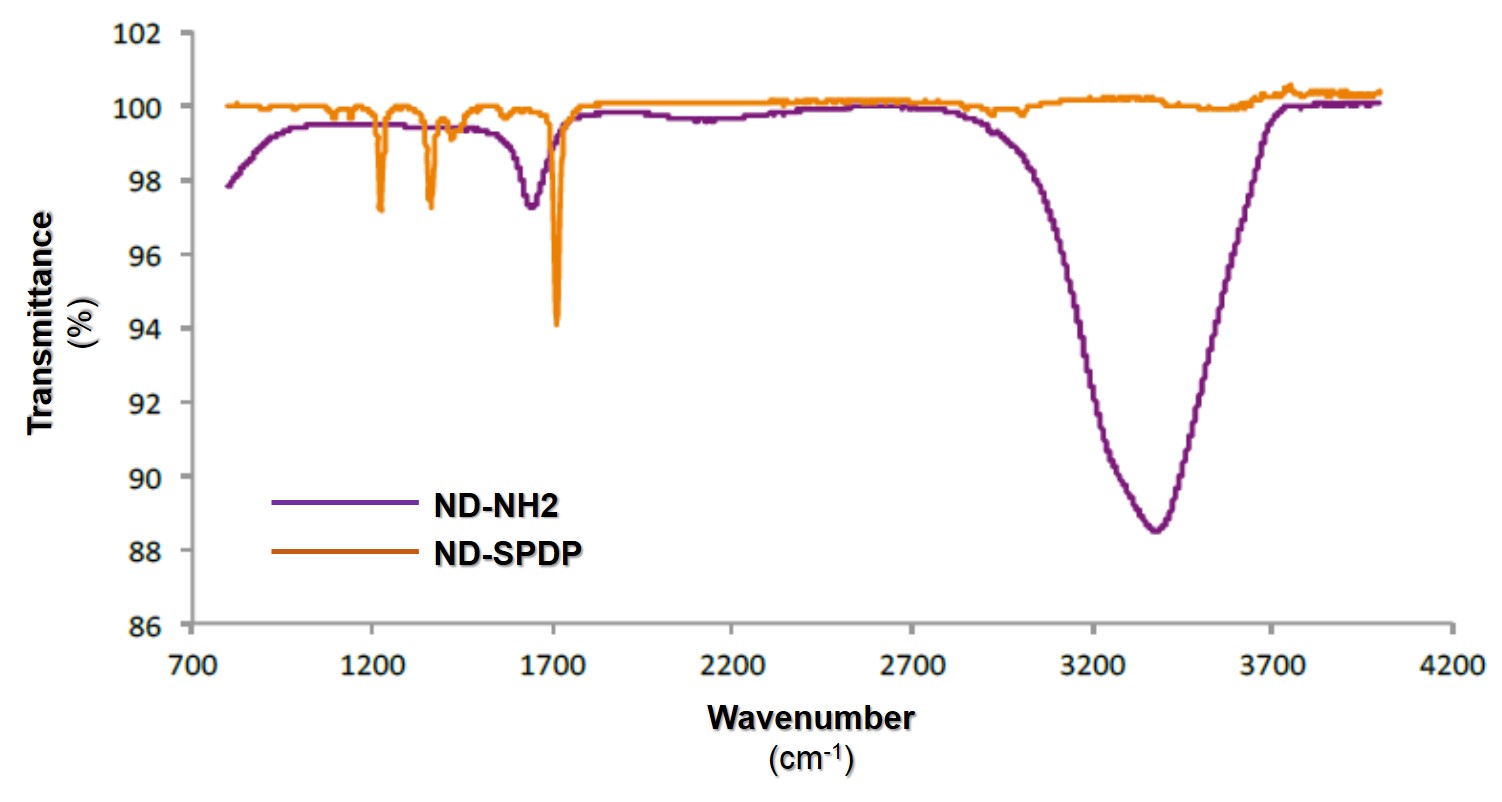

***Supplementary Figure 17***

Confirmation of uptake of Alexa Fluor® 488-labelled nanodiamonds by human pulmonary epithelial cells using flow cytometry.

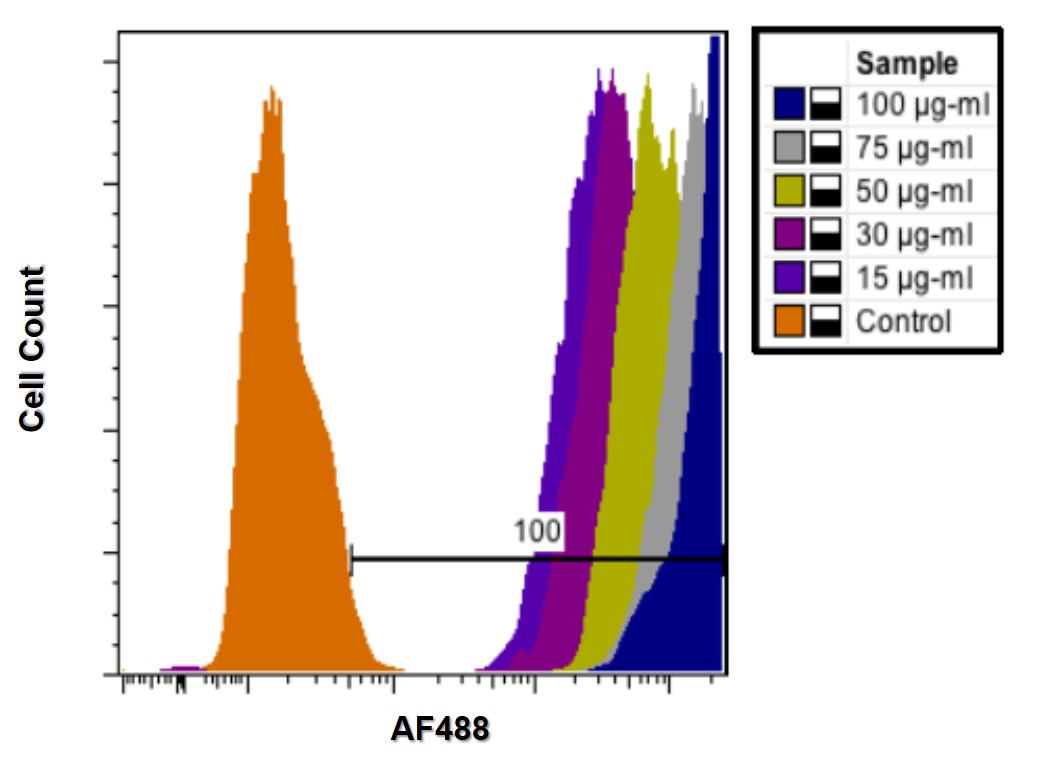
