## Supplementary methods & results for "Prenatal VEGF Nano-Delivery Reverses Congenital Diaphragmatic Hernia-Associated Pulmonary Abnormalities"

***Atomic Force Measurement***

The stiffness of the crosslinked PEG-HCC gel was analyzed after one week of ex vivo lung tissue culture. The crosslinked gel then a fragment of gel was manually removed and stored in a PBS solution for AFM analysis. All the samples were analysed by using an Atomic Force Microscope, mounted on an Inverted Optical Microscope (XEBio, Park Systems, Korea) as previously reported^1^.

***Ex vivo culture of mouse and human fetal lung tissue***

Wild-type CD-1 mice were mated and marked as E0.5 pregnant when they presented a vaginal plug. Pregnant mice were euthanized by cervical dislocation at E12.5, and the embryos were extracted. Lungs were dissected under a dissecting microscope. The trachea was divided just below the larynx. Lungs were washed once in PBS and embedded in 10μL of 100% Matrigel Growth Factor Reduced (MRF) solution (Corning) on top a cell culture insert Transwell® membrane (Corning). After Matrigel gelation, 500 μL of DMEM/F12 medium (Thermo Fisher Scientific) supplemented with 1% p/s (Thermo Fisher Scientific) and 0.1% bovine serum albumin, BSA (Sigma-Aldrich) was added. Culture medium was replaced every other day.

Due to the larger dimensions of human compared to mouse lungs at this stage, we isolated 1x1mm cubes of distal lung, comprising distal epithelium and mesenchyme, and individually embedded them into Matrigel droplets. We selected distal tissue as our models showed that these regions are more affected by mechanical stress (Supplementary Figure 2D).

Human fetal lung samples aged 7 – 8 pcw were dissected into small distal pieces. Each fragment was washed once in PBS and embedded in 10 μL Matrigel (Corning, 356231) on a transwell membrane. After Matrigel gelation, 500 μL basic cell culture medium (DMEM/F12 supplemented with 1% p/s (ThermoFisher Scientific) and 10% FBS (ThermoFisher Scientific) was added per well. Fragments were cultured for 7 days at 37°C, 5% CO_2_, and imaged at 24-hour intervals using a Leica DMi8 microscope with ZEN software. Fresh medium was replaced every other day. For VEGF studies fragments were cultured as described above for the first week and then for one additional week by supplementing 50 ng/mL human recombinant VEGFA (R&D Systems) or 1μM VEGF inhibitor SU5416 (Sigma-Aldrich).

***Haematoxylin & Eosin (H&E) staining of fetal mouse lungs cultured ex vivo***

Mouse lungs cultured *ex vivo* with or without mechanical confinement were fixed in 4% paraformaldehyde (Sigma-Aldrich) for 2 hours at 4°C, washed twice with PBS, incubated with 10 nM NH_4_Cl for 1 hour in and washed. Afterwards, they were dehydrated with sucrose 30% overnight, included in OCT (Optimal Cutting Temperature) compound (Thermo Scientific), cooled in dry ice and sectioned at 7 μm with a cryostat (Bright Instruments).

For H&E staining, sections were hydrated through decreasing grades of ethanol (100%, 100%, 90% and 70%). Then, they were stained in Harris Hematoxylin (Richard Allen Scientific) for 3-5 minutes and rinsed in running tap water for 3 minutes. Sections were differentiated briefly in 1% acid alcohol (70% ethanol + HCl) and rinsed with tap water for 2 minutes. They were counterstained in 1% eosin (Microbiology) for 5 minutes and rinsed in running tap water. Then, the sections were dehydrated through increasing grades of ethanol: 70% (one dip), 90% (one dip), 100% (1 minute) and 100% (1 minute). Finally, they were cleared in Xylene (Honeywell) for 6 minutes and mounted in Dibutylphthalate Polystyrene Xylene (DPX) mounting media (Sigma-Aldrich). Sections’ images were acquired with a NanoZoomer S60 C13210 slide scanner (Hamamatsu).

***Bulk RNA-sequencing and bioinformatic analysis***

Total RNA was isolated with the RNeasy Micro kit (Qiagen), according to manufacturer’s instructions. Reverse transcription to cDNA was performed using the high-capacity cDNA reverse transcription kit (Thermo Fisher Scientific), according to manufacturer’s instructions. Libraries were generated with a KAPA mRNA HyperPrep kit (Roche). Libraries were sequenced on a Novaseq 6000 SP-100 machine (Illumina).

Bioinformatic analysis was performed in R v4.2.2. Genes were filtered out if not having at least 3 replicates from a single condition with at least 2 counts. A variance stabilizing transformation (VST) was performed before Principal Component Analysis (PCA), both performed by the R package DESeq2 v1.38.3^2^. Differentially expressed genes (DEGs) between Day 2 control and Day 2 compression conditions were computed using DESeq2 starting from raw count data, after *parametric fitType*, using a p-value, corrected by Benjamini Hochberg method, lower than 0.05. Results are reported in Supplementary Figure 1. Volcano plot was generated using R package *ggplot2* v3.4.1. A Gene Set Enrichment Analysis (GSEA) within Gene Ontology-Biological Processes database (GO, <https://www.ebi.ac.uk/GOA>) was performed using the R package ClusterProfiler^3^ using a p-value, corrected by Benjamini Hochberg method, lower than 0.01. The results were input in ClueGO v2.5^4^ within Cytoscape v3.9^5^ environment for category clustering (according to score) and network visualization.

***Single-cell dissociation of human fetal lung fragments***

Cultured fetal lung fragments were extracted from Matrigel (Corning, 356231) after 7 days. 2 – 4 control and model samples were pooled respectively into 15 mL conical tubes containing 2 mL Cell Recovery Solution (Corning, 354253), and incubated for 30 min on ice with gentle distressing by inversion every 10 min. Samples were then centrifuged at 500g for 5 min at 4°C, and the supernatant aspirated. Fragments were resuspended in enzyme mixes from MACS Neural Tissue Dissociation Kit P (Miltenyi Biotec, 130-092-628) according to manufacturer’s instructions, and gently triturated 5 times every 5 minutes firstly using a 21G needle (after 5, 10, 15 min cumulatively), and then a 23G needle (after 20, 25, and 30 min cumulatively). The single-cell suspensions were diluted up to 10 mL with wash buffer (1% BSA/HBSS), centrifuged as before, and resuspended in approx. 200 μL resuspension buffer (0.05% BSA/HBSS). Total cell count and viability were assessed for both control and model conditions, and any remaining cell clumps were filtered using a 40 μm Flowmi^TM^ cell strainer. Finally, the cell concentrations were adjusted as required for loading onto the 10X platform.

***Single-cell RNA sequencing and bioinformatic analysis***

Following dissociation, samples were processed for 5’ single-cell capture according to manufacturer’s instructions using 10X Chromium Next GEM Single Cell V(D)J Reagent Kits V2.0 (Rev A Guide). Cells were loaded at a concentration of 700 – 1200 cells/μL with a target recovery of 5000 cells. cDNA from each sample was cleaned up and amplified according to manufacturer’s instructions. Quantification and quality of cDNA was determined using a high-sensitive DNA bioanalyzer. Samples were sequenced by UCL Genomics using a NovaSeq 6000 S4 Flowcell, aiming for a minimum of 50,000 PE (paired-end, 150-bp) reads per cell for gene expression libraries. The 10x raw data were processed using Cellranger v6.1.2 with human genome ex-GRCh38-2020-A^6^. Multiple quality control thresholds were applied; cells with fewer than 250 expressed genes, a mitochondrial gene expression >20%, or a log-transformed UMI count of < 0.80 per cell were removed. Additionally, genes expressed in <0.1% of cells were excluded form analysis. Gene expression values for each cell were then normalized using the global-scaling normalization method ‘LogNormalize’. To mitigate the effects of cell cycle heterogeneity in our dataset, the variations in gene expression associated with G2M/S phases were regressed using the method provided by Seurat v4.2^7^. Principal component analysis was performed using the top 2000 highly variable genes; the top 30 principal components were then selected as input for the harmony algorithm to correct for batch effects between donors and compute a batch-corrected graph^8^. Leiden clustering was performed on this graph and the resolution which best separated all refined cell types, with known differentially expressed markers, was selected. Subclusters were manually examined and further reclustered when necessary. Cell markers were identified in each cluster using the function ‘FindMarkers’, with a 25% minimum expression percentage threshold. UMAP illustration and graphical visualisation of cell type proportions across samples were generated using the ShinyCell application^9^. DESeq2 was used to perform pseudobulk differential expression analysis on specific cell type clusters^2^. Briefly, the raw counts after QC filtering of cells were extracted from Seurat where the counts and metadata aggregated to the sample level. The ‘DESeqDataSetFromMatrix’ function was used to transform cell type cluster counts, metadata and groups into a DESeq data format as a preparation step for differential expression. The negative binomial Wald statistics was used to identify differently expressed genes between model and control. Gene ontology analysis was performed on gene DE genes (FDR < 0.05) using R/Bioconductor cluster Profiler^3^, plotting the top 10 terms for each group or cell type cluster. The epithelial cell compartment was isolated using expression of standard epithelial markers. Epithelial tip cells were further identified by co-expression of SOX9 and TESC genes. The relative proliferative signature, defined by the expression of PBK, BIRC5, MKI67, UBE2C, TOP2A, TK1, AURKB, CDKN3, CENPF, CDK1, and ZWINT genes (from Travaglini et al 2020^10^), was assessed between compressed and control tip cells.

**FEM analysis**

We reconstructed the geometries of the lung, PEG-HCC confinement, and surrounding Matrigel. We assumed an almost-incompressible, isotropic, and almost-linear elastic behaviour for the two materials, adopting a neo-Hookean hyperelastic model. We set the initial Young’s modulus to 18.5 kPa for PEG-HCC and 0.5 kPa for Matrigel (Figure 2B). For the lung tissue we assumed an isotropic and linear elastic constitutive model with Young’s modulus 1 kPa and Poisson’s ratio 0.1, corresponding to a high volumetric compressibility. The proliferation of the lung tissue was modelled assuming a simple law, assuming a transversally isotropic increase of the volume over time.

***Whole mount immunostaining of* ex vivo *culture of mouse and human fetal lung***

EdU assays (ThermoFisher Scientific) were performed according to manufacturer’s instructions and combined with immunofluorescence analysis. EdU incorporation was performed by adding EdU to culture medium for 30 min (mouse fetal lungs) or 1h (human fetal lungs) before fixation. Mouse fetal lungs and human fetal lung samples were mechanically removed from the culture insert and treated with Cell Recovery solution (Corning) for 30 min at 4°C to dissolve Matrigel. Matrigel-free samples were washed once with PBS and fixed in 4% PFA for 30 min at 4°C with gentle rocking. Samples were then dehydrated by washing them once in 25% methanol/PBS (v/v), once in 50% methanol/PBS, once in 75% methanol/PBS, and twice in 100% methanol. Dehydrated samples were stored at -20°C in 100% methanol and used within a few weeks. For rehydration samples were washed once in 75% methanol/PBST (PBST: 0.1% Triton-X in PBS), once in 50% methanol/PBST, once in 25% methanol/PBST and twice in PBST. Samples were treated with 100 mM NH_4_Cl in PBST 0.1% for 1h at room temperature with gentle rocking and incubated in blocking solution (5% Horse Serum in 0.5% PBST) for 1h at room temperature with gentle rocking. Anti-E-Cadherin primary antibody (R&D Systems AF648) was diluted 1:200 in blocking solution and incubated for 36h at 4°C. Samples were washed 5 times with PBST 0.5% overday. Secondary antibody were diluted in blocking solution and incubated overnight at 4°C. Samples were washed 5 times with PBST 0.5% and mounted with 2,20-thiodiethanol, TDE (Sigma-Aldrich). Images were acquired under a Zeiss LSM 710 confocal microscope.

***Immunohistochemistry***

Healthy fetal lung tissue and post-mortem CDH fetal lung tissue were obtained from the GOSH tissue bank at Great Ormond Street Hospital under the REC code 18/EE/0150. For human fetal lung tissue sections, immunohistochemistry analysis was performed through a Bond-Max instrument (Leica). Anti-Ki-67 clone K2 primary antibody (PA0230, pre-diluted) was used in combination with Leica Heat-induced epitope retrieval (HIER) Bond Epitope Retrieval Solution 2 at high pH for 20min, followed by Bond Polymer Refine Detection (DAB), that contains a peroxide block, post primary, polymer reagent, DAB chromogen and hematoxylin counterstain and DAB Enhancer step. Anti-TTF-1 clone SPT24 primary antibody (PA0364, pre-diluted) was used in combination with HIER Bond Epitope Retrieval Solution 1 at low pH for 30min, followed by DAB and DAB Enhancer step (all from Leica). Anti-Flk1 primary antibody (Santa Cruz sc-6251, dilution 1:100) was used in combination with HIER with Leica Bond Epitope Retrieval Solution 2 at high pH for 20min, followed by Bond Polymer Refine Red. Images were scanned with a NanoZoomer S60 C13210 slide scanner (Hamamatsu) and processed with ImageJ software. Briefly, epithelial buds were manually selected and marked as a Region of Interest (ROI). Colour deconvolution was applied to isolate the chromogen signal and the nuclear counterstain. Expression of VEGF and KDR was quantified in each ROI as the mean grey intensity and normalized for the number of nuclei in the same ROI. The percentage of Ki67 and NKX2-1-positive cells was calculated as the number of positive cells divided by the number of nuclei in the same ROI.

For in vivo animal experiments, paraffin sections were placed on polylysine coated slides. All processing was performed with a DAKO Autostainer/Autostainer Plus instrument using DAKO reagents. Pre-treatment deparaffinization rehydration and heat induced epitope retrieval was performed using a 3 in 1 procedure with Envision FLEX target retrieval solutions and system methods. Primary antibodies were used according to the instructions provided by the antibody provider. Secondary antibody and staining were performed using the DACO Envision FLEX/HRP solutions and method programme for the instrument.  This system gives a positive labelling of brown granules in the positive locations within cells or membranes. Thrombomodulin (CD31; platelet endothelial cell adhesion molecule 1, PECAM-1) was used as an endothelial marker/surrogate measure of vascularisation [rabbit anti-thrombomodulin polyclonal antibody (LS-C744612); LifeSpan BioSciences Inc; cross reactivity rat, mouse and human; dilution 1:500 with a positive control of rat placenta. Surfactant Protein-C (SP-C) was used as a marker of type II alveolar epithelial cells/surrogate measure of alveolar epithelial maturation [rabbit polyclonal antibody (FL-197); Santa Cruz Biotechnology Inc; cross reactivity rat and human; dilution 1:100 with positive controls of adult and 3-month old rat lung tissue. Stained sections were imaged using a whole slide scanner (Axioscan Z; Carl Zeiss, Oberkochen, Germany) then rescaled to 25%. A Matlab algorithm (v13a, Mathworks Inc, Natik, USA) allowed the selection of 25 random areas of the lung that were 400 x 400 pixels in size. Color deconvolution was performed to separate the brown and the blue regions of the images^11^. We trained the algorithm to automatically detect SPC-expressing type II pulmonary epithelial cells or thrombomodulin expressing endothelial cells and calculate the SPC+ or thrombomodulin+ immunohistochemistry (IHC) Index (based on the ratio of the number of SPC+/thrombomodulin+ cells over total cell number in pre-determined areas of lung parenchyma).

***Primary Human Pulmonary Epithelial Cell Isolation and Culture***

Epithelial cells were isolated from fresh human lung airway tissue using an enzymatic dissociation method. Briefly, bronchi were cut into 5mm pieces and incubated in a pronase solution (0.15% in DMEM, w:v, Roche Switzerland) overnight at 4^o^C with gentle agitation. Cells were then collected by centrifugation and seeded into 75cm^2^ tissue culture flasks at a density of 10^6^ cells per 25cm^2^ in bronchial epithelial cell growth medium (Lonza, Switzerland). Cells were cultured at 37^o^C and 5% CO_2_ and media were refreshed every 48 hours until they were passaged at day 14 of culture. For cell maintenance, cells were seeded into 75cm^2^ flasks at a density of 3,500 cells per cm^2^.  For all pulmonary epithelial cell experiments cells were seeded in plates/chambers slides pre-coated with human type I collagen (Vitrocol, Advanced Biomatrix, USA) according to manufacturer’s instructions.

***Production of Nanodiamond-Fluorophore Conjugates***

Nanodiamonds (ND) with a primary amine group (ND-NH2) were chemically linked to Alexa Fluor® (AF) 488 via a carboxylic acid succinimidyl-ester. The N-hydroxysuccinimide (NHS) ester is highly reactive towards primary amine groups and results in a stable amide bond (Supplementary Figure 12). The Alexa Fluor ® 488 succcinimidyl ester was linked to the ND-NH2 particles at a weight ratio of 2:1, in a 50:50 volume mix of DMSO and 0.1 M sodium bicarbonate buffer (pH=8). The reaction was maintained under stirring conditions for 3 hours, protected from light, at room temperature. The solution was then washed 5 times with acetone and twice with ddH2O via repeated centrifugation cycles (14.8K RPM, 90 minutes, 4°C). The resulting solution was subsequently transferred to a pre-weighed Eppendorf tube (Eppendorf, Hamburg, Germany) and placed in a SAVANT DNA 120 speedvac concentrator (Thermo-Scientific, USA) to dry overnight. The AF conjugated ND (ND-AF) yield was calculated by subtracting the net weight of the empty Eppendorf tube.

***Measurement of Fluorophore Amount on Nanodiamond-Fluorophore Conjugates***

AF dye conjugation and the amount of dye conjugated to the ND surface were determined by ultraviolet-visible spectroscopy (UV/Vis) using a Varian Cary 300 Bio instrument and the Cary WinUV software package (Agilent, USA). The AF and ND-AF solutions were diluted 1/1,000 in ddH2O and their absorbance measured at 494nm and at 650nm for AF488 and AF647 dyes respectively. The dye concentration was determined using the Beer-Lambert equation: A=ε.l.c, where A= absorbance (a.u.), ε= molar extinction coefficient (M-1cm-1), l= path length (cm), c= concentration (M); given a path length of 1 cm and molar extinction coefficient of 71000 M-1cm-1. Concentrations were multiplied by the dilution factor and subsequently the degree of AF labelling on the ND surface was calculated using the following equation: (concentration of dye on ND surface / concentration of free dye) x 100%.

***In Vitro Nanodiamond Cellular Uptake and Toxicity***

Human pulmonary epithelial cells were co-cultured with increasing concentrations (5-100μg/mL) of ND (ND-AF488: cellular uptake and localisation; ND-NH2: cytotoxicity and proliferation) for up to 72 hours. The degree of cellular uptake (proportion of cells positive for AF488 expressed as a percentage of total, as well as log degree fluorescence) was assessed by FACS analysis, and localisation was assessed by confocal microscopy. For FACS analysis, cells were seeded in 24-well plates in duplicate at a density of 60,000 cells per well; the following day cells were incubated with ND-AF488 at the desired concentrations for 4 hours at 37oC, washed and analysed using a LSR Fortessa instrument (BD Biosciences, UK). For confocal microscopy, we performed live cell imaging of cells plated in 8-well chamber slides (Ibidi, Germany) at a seeding density of 15,000 cells per well; after allowing time for attachment, cells were incubated with ND-AF488 at the desired concentration for up to 72 hours, and subsequently washed and stained with Hoechst33258 (5μg/mL in media; Hoechst, Germany) prior to confocal imaging using an inverted LSM700 confocal laser-scanning microscope (Zeiss, Germany).

To assess cytotoxicity, pulmonary epithelial cells (duplicates; 10,000 cells per well; 96-well plate) incubated with ND-NH2 for 72 hours were stained with Propidium Iodide (PI; 50μL of 3μg/mL solution in PBS, Invitrogen, USA,) and Annexin V (FITC Annexin V, 1/100 in binding buffer, v:v, Biolegend, USA;) as described previously. On FACS analysis (performed on a FACSAria instrument, BD Biosciences, UK), Annexin V-/PI- cells were considered to be alive, Annexin V+/PI- in early apoptosis, Annexin V+/PI+ in late apoptosis, and Annexin V-/PI+ cells were considered to be dead. To test cellular metabolic activity, we used the well-established Alamar Blue assay. Briefly, cells were seeded in triplicate (10,000 cells per well; 96-well plate), and after attachment were incubated with ND-NH2 for up to 72 hours. At the chosen analysis timepoints (24, 48, 72 hours), Alamar Blue (1/10 in media, v:v, Sigma, USA) was added and absorbance values at 570nm were read with a Titertek Multiscan MCC/340 plate-reader (Labsystems, Finland) and normalised to absorbance values at 600nm.

5-ethynyl-2’-deoxyuridine (EdU; click-iT EdU-Pacific Blue Flow Cytometry Kit, Invitrogen, USA; FACS analysis) and bromodeoxyuridine (BrdU; 1/100 in cell culture media, v:v, Invitrogen, USA; confocal microscopy) assays were used to assess cellular proliferation of pulmonary epithelial cells incubated with increasing concentrations of ND-NH2 for 72 hours. For EdU assays, cells were plated in 24-well plates at a density of 50,000 cells per well and following attachment were incubated with ND-NH2 for 48 hours. The EdU kit was used in accordance with the manufacturer’s protocol with an EdU (10μM) incubation time of 14 hours, and analysis was performed on an LSRII instrument (BD Biosciences, UK). For BrdU assays, cells were grown in chamber slides (Ibidi, Germany) at a density of 15,000 cells pre well and after 48 hours of treatment with ND-NH2 cells were incubated with BrdU labelling agent (1/100 in media, v:v, Invitrogen, USA) overnight. After fixation (70% ethanol for 30 minutes), denaturation of genomic DNA (1.5M hydrochloric acid for 30 minutes) and blocking (5% normal goat serum, 0.3% TritonX-100 in PBS), primary rat anti-BrdU IgG2a antibody was added to chamber slides (1/200 in blocking solution, v:v, AbD-Serotec, UK) and incubated overnight at 4oC. The next day cells were incubated with AF633 goat anti-rat secondary antibody (1/300 in blocking solution, v:v. Invitrogen, USA) for 3 hours at room temperature. Cells were washed an immersed in PBS prior to confocal imaging with a 63X oil immersion lens, a Helium Neon laser excitation of 633nm and a diode laser excitation of 405nm for AF633 and DAPI detection respectively. Cells treated with secondary antibody alone served as negative controls. In order to visualize cell boundaries and potential cytoskeletal insults resulting from ND treatment, BRDU stained cells were also incubated with anti-cytokeratin rabbit polyclonal IgG primary antibody (1/200 in blocking solution, v:v; Abcam, UK) and labelled with Alexa Fluor® 488 donkey anti-rabbit IgG H+L (1/300 in PBS, v:v; Invitrogen, USA,) secondary antibody.

***Recombinant Vascular Endothelial Growth Factor Production and Purification***

Recombinant vascular endothelial growth factor-A (VEGF-A; VEGF164) production was carried out using Pichia pastoris yeasts from the laboratory of Prof. P. Carmeliet (Vesalius Research Center, KU Leuven, Belgium) as described before. VEGF was purified from the cell supernatant by affinity chromatography, with heparin as ligand, followed by size exclusion chromatography. Purity and integrity of VEGF were checked by SDS-Page and concentration was evaluated by UV spectrophotometry (calculated dimer molar extinction coefficient: e = 12,115 M^−1^ cm^−1^) and ELISA. The VEGF solution was filter-sterilized, and its concentration adjusted to 1 mg/mL. The activity of VEGF was assessed by detection of VEGF-receptor-2 (KDR/Flk1) phosphorylation of human umbilical vein endothelial cells (HUVEC) cells by Western blot.

***Assessment of Morphology, Size, and Surface Charge of Nanodiamond-VEGF Conjugates***

Transmission electron microscopy (TEM) was performed to ascertain ND-NH_2_ and ND-VEGF morphology. ND-NH_2_ and ND-VEGF solutions were directly drop-caster and dried on 200-mesh copper grids (manufactured by Dr. Mark Tumaine, Department of Anatomy and Developmental Biology, UCL). ND-NH_2_ and ND-VEGF were then viewed with a Jeol 1010 TE Microscope and the images recorded using a Gatan Orius CCD Camera (Gatan, UK)

Size, size distribution and surface charge of ND-VEGF (as well as ND-AF488 and ND-NH_2_) were determined by dynamic light scattering (DLS) and by surface potential (ζ potential) measurements using a Zetasizer NanoZS instrument (Malvern, UK). All measurements were performed at 25^o^C and at 173^o^ scattering angle. Cumulative analysis of 90 measurements (20 seconds per run) was performed and the percentage number distribution was estimated. Output measurements included mean diameter and polydispersity index (PDI),

***Production of Nanodiamond-VEGF Conjugates***

The process of ND-VEGF production is summarised in Supplementary Figure 11A. ND-NH_2_ particles were first coupled to a sulfhydryl-to-amine cross-linker Sulfosuccinimidyl 6-(3'-[2-pyridyldithio]-propionamido) hexanoate (Sulfo-LC-SPDP), which contains an amine-reactive NHS ester group and a thiol-reactive pyridyldithiol group. The NHS ester arm of the linker reacts with amine terminated nanodiamond particles to form a stable amide bond, while the pyridyldithiol arm reacts with thiol-modified (thiolated) VEGF to yield reducible disulphide bonds^12,13,14^ (Supplementary Figure 11A). The Sulfo-LC-SPDP (Thermo-scientific, Waltham, MA, USA) was coupled to ND-NH_2_ particles overnight, at a molar excess of 377 in a 0.1 M sodium bicarbonate buffer (pH 8). The reaction was performed under stirring conditions and at room temperature. The resulting solution was then washed thrice with ddH_2_O via repeated centrifugation cycles (14.8K RPM, 90 minutes, 4°C), and subsequently transferred to a pre-weighed Eppendorf tube and placed in a SAVANT DNA 120 speedvac concentrator (Thermo-Scientific), to dry overnight. ND-SPDP yield was calculated by subtracting the net weight of the empty Eppendorf. The subsequent linking of ND-SPDP with VEGF made use of 2-Iminothiolane.HCL (Traut’s reagent, Thermo-Scientific), a thiolation compound that reacts with primary amines to yield accessible sulfhydryl groups^15,16^. The Traut’s reagent was linked with lyophilized rat VEGF at a molar ratio of 4:1 in 0.1 M sodium bicarbonate buffer (pH 8), for 4 hours under stirring conditions, and at room temperature.  The resultant mixture was then filtered through Illustra NAP-10 columns (GE Healthcare Life Sciences, Amersham, UK) to permit solvent exchange of residual PBS with sodium bicarbonate buffer. To quantify the yield of VEGF resulting from the thiolation reaction, a BCA assay (Thermo-Scientific) was performed according to the manufacturer’s instructions.  In the final step the ND-SPDP functionalized particles were linked to the thiolated-VEGF at a molar ratio of 1:1. The reaction was performed in 0.1 M sodium bicarbonate buffer (pH 8) overnight under stirring conditions, and at room temperature. The following day, the resulting product was washed thrice with ddH_2_O, via sequential ultracentrifugation cycles (45K RPM, 90 mins, 4°C), and resuspended in 1 mL of ddH_2_O.

***Measurement of VEGF Amount on Nanodiamond-VEGF Conjugates***

To determine the quantity of VEGF conjugated to ND we used a rat VEGF ELISA kit according to manufacturer’s instructions (100786, Abcam, Cambridge, UK). Briefly, 100 µL of ND-VEGF samples of various dilutions and VEGF standards supplied with the kit were pipetted into a 96-well plate, pre-coated with rat VEGF antibody, in duplicate and incubated at 4°C overnight on a horizontal orbital microplate shaker at 500 RPM. The plate was washed 4 times with wash buffer, using an automated Denley Cellwash plate washer (Triad Scientific, USA) and 100 µL of biotinylated VEGF detection antibody was added to each well and incubated for 1 hour, with gentle shaking at room temperature.  Following this, the plate was washed 4 times and 100 µL of horseradish peroxidase streptavidin was added to each well for 45 minutes, in order to bind to the biotinylated detection antibody. The plate was subsequently washed 4 times and 100 µL of 3,3',5,5'-Tetramethylbenzidine (TMB) substrate (chromogenic solution that develops proportionally to the amount of VEGF bound) was added to each well and incubated for 30 minutes, protected from light with gentle shaking. 50µL of stop solution was then added to quench the substrate reaction and the plate absorbance was read at 450 nm, using a Titertek Multiscan MCC/340 plate reader (Labsystems, Vantaa, Finland). Absorbance values from samples were compared to a generated standard curve to ascertain the concentration of VEGF on ND-VEGF.

***Assessment of Thiolated-VEGF Biological Activity***

In order to ensure that the thiolation process does not affect the biological activity of VEGF used in this series of experiments, we utilized the well-established chorioallantoic membrane (CAM) assay (42). Fertilized chicken eggs (Henry Stewart and Co, UK) were incubated at 37°C. At day 3 of incubation an oval window of approximately 3 cm in diameter was cut into the shell with small dissecting scissors (World Precision Instruments, UK) to reveal the embryo and CAM vessels were sealed with tape. At day 8 of incubation 2 mm diameter filter papers were either soaked in PBS (negative control), 100ng/mL unmodified VEGF (positive control) or 100ng/mL thiolated VEGF and placed on the CAM. Samples were examined and photographed *in ovo* with a stereomicroscope equipped with a camera system up to day 18 of incubation. The number of blood vessels less than 10 μm in diameter converging towards the placed papers were counted blinded reporting the mean of five counts.

***VEGF Release Profile from Nanodiamond-VEGF Conjugates***

In order to assess the release profile of VEGF conjugated on ND, dithiothreitol (DTT; Sigma-Aldrich; 10mM in ddH_2_O, pH 7) was added in ND-VEGF solution (final concentration of DTT 1mM). VEGF desorption was tracked 6-hourly for 72 hours. Individual samples of ND-VEGF (with and without DTT; from the same original solution) were prepared for each timepoint and kept at 37^o^C.  Once a timepoint was reached, allocated samples were ultracentrifuged (45K RPM, 90 mins, 4°C) to pellet ND, and the supernatant was collected for analysis. Supernatant VEGF concentration was assessed by ELISA as described above. The percent of VEGF release/desorption was based on the amount of VEGF initially conjugated on ND and on the amount of free VEGF in the supernatant. The experiment was performed in triplicate.

***Surgical procedures***

On E19, rats were sedated and anesthetized with 2% isoflurane (Isoba Vet, Abbott Laboratories Ltd, Queensborough, Kent, UK) and placed on a heated pad (37 °C). The uterus was exposed by a midline laparotomy. The fetuses were counted and examined under micro-ultrasound using a MS400 (18-38 MHz) probe on a Vevo 2100 Imaging System (Visual Sonics, Toronto, Ontario, Canada). When the presence of a diaphragmatic defect was confirmed, 3-4 of such fetuses per rat were either used in a preliminary *“biodistribution study”* (see also Supplementary Results and Supplementary Figures 12,13 for details) or were randomized to an experimental group of the main *“intervention study”* using an online randomization tool (see below for experimental group details). Tracheal occlusion and tracheal injections were performed as previously described^17,18^. Briefly, a 6-0 polypropylene (Prolene, Ethicon, Zaventem, Belgium) purse-string suture was placed over the fetal head area, where a hysterotomy of 0.8 cm was performed. The head and neck were exposed, and the purse-string was tightened to keep the rest of the fetus inside the uterus. The fetal head was hyperextended and, under stereoscopic zoom microscopy (x10 magnification), the fetal trachea was exposed by sharp and blunt dissection. Under direct vision, tracheal injections were performed using a 50μL Hamilton Glass Syringe with a 30 G sharp needle (Hamilton, Bonaduz, Switzerland). After the injections the trachea was ligated using a micro titanium clip (Weck Horizon, Teleflex, Wayne, USA). Finally, the head was internalized into the uterus and the purse-string suture closed after the injection of 0.5-1 mL of warm saline isotonic solution. The uterus was then placed in the abdomen, closed in layers with 5-0 polyglactin 910 (Vycril, Ethicon) running suture. The incision was infiltrated with 2% lidocaine hydrochloride (Linisol: B Braun) and buprenorphine 0.05mg/kg (Temgesic: Schering-Plough, Brussels, Belgium) was injected SC for pain relief. Rats were kept on a heating pad (37°C) until they are fully recovered. On E21, dams were anesthetized with 2% isoflurane (Isoba Vet: Abbott) and euthanized by cervical dislocation. Fetuses were harvested when movements were no longer identified. Fetal body weight was recorded. The abdominal cavity was opened under stereoscopic zoom microscopy (10x magnification) to confirm the CDH. Lungs and other tissues were harvested, weighed and either placed in O.C.T. compound (Sakura, Finetek, USA) and frozen in liquid nitrogen, or fixed in 4% paraformaldehyde.

***Experimental groups for* in vivo *intervention study***

A total of 178 fetuses (10 healthy, 168 with CDH) from 55 time-dated pregnant rats were included in the study. The incidence of nitrofen-induced CDH in the present series of experiments was 40%. There were eight groups in the main *in vivo* intervention study (summarised in Figure 4A): (i.) healthy control (non-CDH; olive oil gavage-fed mothers), (ii.) sham intervention (surgery but no injection or TO), (iii.) CDH+PBS+TO (50μl of PBS per fetus), (iv.) CDH+VEGF+TO (100ng free VEGF per fetus), (v.) CDH+ND+TO (12μg ND-NH_2_ per fetus), (vi.) CDH+ND-VEGF+TO (100ng ND-conjugated VEGF per fetus), (vii.) CDH+SU5416+TO (6μg of the KDR/Flk1 inhibitor SU5416 per fetus) and (viii.) CDH+ND-VEGF+SU5416+TO (100ng ND-conjugated VEGF and 6μg of SU5416 per fetus). Animals in the groups with ND-VEGF administration (ND-VEGF+TO, ND-VEGF+SU5416+TO) received 13.4±1.9μg ND and 100ng VEGF (concentration in injection solution of 267.8±39.5μg/mL and 2μg/mL respectively), while animals in ND+TO and VEGF+TO received the same amounts of empty ND (ND-NH_2_) and free VEGF respectively, in 50μL PBS. The concentration of the KDR/Flk1 inhibitor SU5416 in the injection solution used in SU5416+TO and ND-VEGF+SU5416+TO groups was 120μg/mL. The investigators were blinded to group allocation during all subsequent procedures and outcome assessments in the intervention study.

***Assessment of survival and toxicity***

Fetal survival was assessed at E21; macroscopically normal pink looking pups with spontaneous limb movements on delivery were considered alive. Survival rate was expressed as the percentage of injected animals found to be alive on harvest. Autopsies were performed by an independent veterinary pathologist (Dr Joy Archer, Queens Veterinary School Hospital, University of Cambridge, UK) blinded to group allocation in tissues (brain, eyes/retina, heart, lungs, liver, spleen, kidney, stomach, intestine) from selected rat mothers, treated CDH fetuses and non CDH littermates to detect any toxic side-effect of the interventions.

***Assessment of lung growth***

The weight of the whole fetus as well as of the lungs (and other harvested tissues were recorded). Lung-to-body weight ratio (LBWR; expressed as a percentage) and lung total protein content were used as surrogate measures of lung growth. Lung protein quantification was done as previously described^19^. In brief, paraffin embedded lung 10 μm thick coronal sections were placed in Eppendorf tubes and deparaffinized in xylene for 10 min, then tissue was pelleted at 12K g for 3 minutes. Both processes were repeated three times. The deparaffinized tissue pellets were rehydrated in ethanol. All samples were then weighed, and matched amounts of tissues were each immersed at a 20% w/v ratio in a 20mM Tris HCl, pH 8.8, 2% SDS, and 200mM DTT extraction buffer. All samples were subjected to high-temperature extraction at 100°C for 20 min, and then at 80°C for 2h with shaking. Extracts were clarified for 15 min at 12 000×*g* at 4°C, quantified by the Bradford method and with the EZQ Protein Quantitation Kit (Molecular Probes, Eugene, OR, USA) and normalized to wet lung weight.

***Lung Morphometry***

The lungs were paraffin embedded, and 4 μm thick coronal sections were taken. The sections were stained with hematoxylin and eosin (H&E) staining for airway morphometry and with Miller’s elastic staining for vascular morphometry. Lung morphometry was done by two operators blinded for treatment allocation using an Axioplan light microscope (Carl Zeiss, Oberkochen, Germany) at 200x or 400x magnification. Measurements were made using two special oculars, one with a grid and the other one with a ruler. Airway morphometry measurements were made on 10 to 20 non-overlapping fields as previously described: the two components of mean linear intercept (Lm) mean wall transection length (Lmw) and mean linear intercept of parenchymal airspace (Lma), as well as mean terminal bronchiolar density (MTBD)^20^ . Vascular morphometry measurements were made on up to 50 peripheral arteries with an external diameter (ED) of 30-50µm in 20 non-overlapping fields. Vascular morphometric parameters included the ED, the internal diameter (ID) and the adventitial diameter (AD), all measured along the shortest axis. From these parameters the following variables were calculated: proportionate medial thickness (%MT = (ED-ID)/ED x 100) and proportionate adventitial thickness (%AT = (AD-ED)/ED x 100)^21^.

**Supplementary Results**

***Nanodiamond In Vitro Characterisation Studies***

In order to test in a validated model our hypothesis we planned to deliver VEGF in Nitrofen rat model of CDH coniugated to nanodiamond (ND) to obtain slow continue release. We firstly designed and validated 2 different ND based platforms. In the first, ND with a primary amine group (ND-NH_2_) were conjugated with the fluorescent dye Alexa Fluor® 488 (AF488; ND-AF488) via a carboxylic acid succinimidyl-ester. The N-hydroxysuccinimide (NHS) ester is highly reactive towards primary amine groups and results in a stable amide bond (Supplementary Figure 12). ND-AF488 were used for in vitro cellular uptake as well as *in vivo* biodistribution studies (Supplementary Figures 13,17). In the second delivery platform ND-NH_2_ were conjugated with recombinant rat or human vascular endothelial growth factor A (VEGF-A; ND-VEGF) using a sulfhydryl-to-amine cross-linker and thiol-modified (thiolated) VEGF (Supplementary Figure 11A). The resulting disulphide bonds could be disrupted under reducing conditions to release biologically active VEGF (Supplementary Figure 11B). The ND-AF488 product used for *in vitro* (cellular uptake and toxicity) and in vivo (biodistribution) studies had a ND concentration of 198.7±6.0μg/mL and AF488 concentration of 154.3±5.0μg/mL (n=3). The loading efficiency of 10μg of AF488 to ND for a 2:1 AF488/ND ratio was 38.9±0.9% relative to the original AF488 mass and 77.7±0.1% relative to the mass of ND added. Fourier transform infrared spectroscopy (FTIR) analysis revealed that AF488 covalently bound to ND-NH_2_ by amide formation (denoted by the appearance of benzene ring stretch signals), and ultraviolet-visible (UV/Vis) spectroscopy confirmed a characteristic peak at 488nm (Supplementary Figure 14). Dynamic light scattering (DLS) analysis demonstrated that ND-NH_2_ form average cluster sizes 47.6±0.2nm and zeta (ζ) potential of 13.8±2.9mV (n=3). Conjugation with AF488 increased the average cluster size by approximately two times (99.2±2.1nm; n=3), suggesting that fluorescent dye conjugation makes ND more prone to agglomeration. ND-AF488 also exhibited increased ζ potential (29.6±1.7mV; n=3), which is indicative of surface binding of the dye on ND. ND-NH_2_ as well as ND-AF488 had a narrow size distribution, with polydispersity indices (PDI) around 0.2. For ND-VEGF utilised in our intervention study, we confirmed successful completion of individual steps in the preparation process using UV/Vis spectroscopy (Supplementary Figure 15). Bonding of the sulfhydryl-to-amine cross-linker Sulfosuccinimidyl 6-(3'-[2-pyridyldithio]-propionamido)hexanoate (Sulfo-LC-SPDP) resulted in a distinct absorbance peak near 270nm (Supplementary Figure 15; also see Supplementary Figure 16 for FTIR confirmation of SPDP coupling), and subsequent disulphide bond formation with (and loading of) thiolated VEGF generated a characteristic VEGF absorbance peak at 209 nm (Supplementary Figure 15). We achieved a ND concentration of 469±8.0μg/mL and a VEGF concentration of 3.5±0.1μg/mL (n=5). The loading efficiency of 10μg of VEGF for a 1:1 VEGF/ND ratio was 1.8±0.1% relative to the original thiolated-VEGF mass and 1.7±0.1% relative to the mass of ND added. ND-VEGF had an average cluster size of 149±3.7nm (as measured by DLS), ζ potential 59.6±2.3mV (consistent with surface binding of VEGF to ND), and PDI of 0.2. Transmission electron microscopy (TEM) of ND-VEGF demonstrated lattice-like structures, which was in contrast to the distinct clusters seen when ND-NH_2_ were imaged (Supplementary Figure 11E). VEGF desorption experiments showed sustained release of VEGF from ND-VEGF over 72 hours in the presence of reducing conditions, which is consistent with gradual disruption of disulphide bonds between VEGF and ND (97±2% of loaded VEGF released within 72 hours; n=3; Supplementary Figure 11B). Using chorioallantoic membrane (CAM) assays, we confirmed that thiolated-VEGF released from ND-VEGF maintained biological activity similar to that of the unmodified/stock compound (number of newly-formed blood vessels after 10 days; VEGF: 24±4 vs. thiolated-VEGF: 25±6; n=3; Supplementary Figure 11C,D). Cellular uptake analysis using human pulmonary epithelial cells showed uptake of ND into the intra-cellular compartment. ND uptake by pulmonary epithelial cells was quick and dose dependent. FACS analysis of cells co-cultured with ND-AF488 demonstrated near complete uptake when a particle concentration ≥75μg/mL was used (Supplementary Figure 11F; also see Supplementary Figure 17) with no significant changes on the percentage of proliferating cells (Edu assays; Supplementary Figure 11G). These findings were confirmed with confocal microscopy of live cells, which also showed heterogeneous cytoplasmic distribution/localisation of ND-AF488 suggestive of endocytosis (Supplementary Figure 11H). The uptake process was completed within 4 hours with no significant changes observed thereafter. The addition of increasing concentrations of ND-NH_2_ to the culture medium of human pulmonary epithelial cells did not significantly alter their metabolic activity (Alamar Blue assay; Supplementary Figure 11I) or affect their proliferation (BRDU assays; Supplementary Figure 11J). Laminin C staining confirmed the absence of any particle-induced cytoskeletal damage (Supplementary Figure 11J). Finally, co-culture of cells with ND-NH_2_ did not result in cytotoxicity; after 72 hours the majority were healthy, with low levels of apoptosis (early/late) and necrosis (PI/Annexin V assay; Supplementary Figure 11K).

***Nanodiamond In Vivo Biodistribution Studies***

The aim of the biodistribution experiments was to determine the localisation of AF488-labelled ND (ND-AF488) in various tissue compartments/organs following in utero intra-tracheal administration in E19 fetal rats following tracheal occlusion (TO). A total of 34 fetuses (all healthy) from 11 time-dated pregnant rats received in utero injections in this preliminary study, and there were four experimental groups: (i.) ND-AF488 with TO, (ii.) ND-AF488 without TO (to determine whether TO is required for uptake of ND in the fetal lung), (iii.) free AF488 with TO, and (iv.) free AF488 without TO. Animals in ND-AF488 groups (with/without TO) received 3.9±0.1μg ND and 3μg AF488 (concentration in injection solution of 77.3±1.8μg/ml and 60μg/mL respectively), while animals in AF488 groups (with/without TO) received the same amount of free AF488 in 50μL of PBS (Supplementary Figure 13A,B). Fetal survival was 65% (9/14) in animals that received ND-AF488 with TO and was similar in all other experimental groups (Supplementary Figure 13C). There was no maternal mortality. Similar to what we observed *in vitro*, *in vivo* uptake of ND-AF488 following intra-tracheal administration at E19 was quick, with fluorescence detected in lung sections from fetuses harvested as early as 24 hours (E20). Uptake increased thereafter and peaked at 48 hours post injection (E21; Supplementary Figure 13D) but was minimal without TO (Supplementary Figure 13E). ND-AF488 were mostly found in distal airways, with evidence of uptake by SPC-expressing type II alveolar epithelial cells (Supplementary Figure 13F). In contrast, there was no AF488 retention at any time-point in the lungs of fetuses with TO injected with free dye (same amount as what was carried on ND-AF488; Supplementary Figure 13G). Examination of other sections from other fetal organs using confocal microscopy demonstrated no particle “spillage” in tissues other than the lung in animals that received ND-AF488 with TO; some particle uptake was seen in the gastrointestinal tract of animals that were given ND-AF488 without TO. ND-AF488 were not detected in the placenta or maternal tissues in any of the experimental groups.
